# Load-Dependent Effects of Sodium Glucose Co-Transporter Inhibitors on Work in Human Hypertrophic Cardiomyopathy Living Myocardial Slices

**DOI:** 10.64898/2025.12.03.692183

**Authors:** Rebecca B. Taichman, Julia N. Smolyak, Jesse Chittams, Kelly Gallagher, Christopher M. McAllister, Jaime M. Yob, Kenneth C. Bedi, Michael P. Morley, Trisha T. Phan, Sapna N. Patel, Shawnaleh Cada, Christopher Petucci, Kenneth B. Margulies, Sharlene M. Day, Benjamin W. Lee

## Abstract

**Background:** Disease modifying therapies for hypertrophic cardiomyopathy (HCM) remain a prevailing unmet need. Human-based experimental platforms capable of controlled manipulation of preload and afterload can distinguish direct myocardial and systemic effects and facilitate development of targeted cardiac therapeutics. Sodium glucose cotransporter inhibitors (SGLTi) may directly affect cardiac contractility, potentially related to increased ketone availability. These effects have not been adequately studied in human HCM myocardium under defined loading conditions.

**Aims:** We sought to establish human living myocardial slices (LMS) as a platform to interrogate load-dependent myocardial mechanics in HCM and to quantify the acute effects of metabolic and pharmacologic interventions—including SGLTi—on myocardial work under physiologic loading conditions.

**Methods:** Human myocardial tissue was procured from non-failing donor hearts or individuals with HCM undergoing septal myectomy. Freshly prepared human LMS were mechanically tested to generate biomimetic work loops across a range of physiologic preloads and afterloads in either glucose-only fuel or glucose supplemented with ketone. Following baseline measurements, slices were loaded with drug (isoproterenol, mavacamten, sotagliflozin, or empagliflozin) or vehicle (DMSO) and work loop analysis was repeated, allowing each slice to serve as its own control. Mixed effects linear regression models incorporating random effects for heart and slice and fixed effects for clinical characteristics evaluated determinants of myocardial work and drug response across loading conditions.

**Results:** A total of 120 LMS from 32 individuals (16 non-failing and 16 HCM) were analyzed. At baseline, myocardial work was positively associated with younger age, hypertension, and ejection fraction. Ketone supplementation augmented work and work-strain slope particularly in HCM LMS at high afterloads. We validated our drug testing methodology by demonstrating increased work with known positive inotrope isoproterenol, decreased work with negative inotrope mavacamtem most pronounced in HCM LMS, and a null effect of DMSO. Acute exposure to SGLTi sotagliflozin and empagliflozin directly reduced myocardial work, with increased potency of sotagliflozin at high afterloads.

**Conclusions:** Our LMS platform enables assessment of myocardial mechanics across controlled loading conditions and is an ideal platform to rigorously phenotype human myocardial tissue and interrogate direct effects of pharmacologic intervention. We demonstrate that SGLTi and ketones have distinct and discordant effects on human myocardial contractility.

## INTRODUCTION

Hypertrophic cardiomyopathy (HCM) is the most common Mendelian cardiovascular disorder, affecting 1 in 500 people.^1,2^ Currently, HCM interventions largely target symptomatic relief of left ventricular outflow tract (LVOT) obstruction.^3^ Multiple highly effective strategies reduce obstructive symptoms, including negative inotropes, invasive septal reduction therapies (septal myectomy or alcohol septal ablation), and myosin modulators like mavacamten and aficamten.^4–6^ However, substantial disease burden remains beyond obstructive symptoms, including impaired diastolic function, atrial fibrillation with embolic stroke, and fatal arrhythmias.^3^ The ODYSSEY-HCM trial demonstrated no therapeutic benefit of mavacamten in non-obstructive HCM,^7^ representing 40-50% of patients with HCM.^8^ Disease-modifying therapy for HCM remains a prevailing unmet need.^3–5,9^

Clinical translation of mechanistic-based HCM therapies remains low, partly due to the limited translation of successful interventions in small animal models.^10–12^ Human-based platforms that rigorously assess drug impact on contractile function under physiologic loading conditions are instrumental for the development and translation of targeted cardiac therapeutics. Human living myocardial slices (hLMS), precision-cut sections of fresh cardiac tissue, are a tool for studying the primary myocardium as a minimum, functional three-dimensional unit, yielding a much higher degree of *in vivo* mimicry.^13–15^ Prior studies of hLMS contractility largely focused on achieving long-term culture^16,17^ however show significant remodeling even during short culture duration^18^ thus limiting their relatability to the antecedent clinical state. Further, prior force measurements largely relied on fixed-length isometric measurements, limiting assessment of load-dependent effects. A standardized heart procurement, slice assessment, and drug testing methodology is essential for reliable drug testing in hLMS, particularly for drugs with both potential systemic and direct cardiac effects, such as sodium glucose co-transporter inhibitors (SGLTi).

SGLTi are remarkably cardioprotective in heart failure, but their direct effect on cardiac contractility has not been fully studied, particularly in the context of HCM. Two potential hemodynamic effects of SGLTi confound assessment of their direct impact on contractility.^19^ First, naturiesis may reduce preload on the heart.^20,21^ Second, a modest systemic ketosis may reduce afterload and/or increase contractility.^22–27^ Studies of ketone supplementation^25,28–31^ confirmed dose-dependent increase in cardiac output coupled with marked vasodilation, raising the question of whether exogenous ketones improve cardiac function through *direct* increase in myocardial contractility, *indirect* reduction in afterload, or a combination of both. SGLT2 is not expressed in the heart and SGLTi confer myocardial protection in mice despite SGLT2 knockout, prompting consideration of other direct myocardial targets.^32–34^

Prior studies examining the acute direct effects of SGLTi on human cardiac tissue mechanics demonstrated inconsistent effects on forces and/or contractile kinetics.^35,36^ With a variety of human heart preparations, including trabeculae, living myocardial slices, isolated cardiomyocytes, and human engineered heart tissues, separate groups found either reduced, null, or increased active contractility as well as a range of effects on diastolic forces and contraction/relaxation times.^35–37^ These inconsistent results indicate that the effect of SGLTi on contractility may be dependent on underlying disease, types of *ex vivo* preparation, or mechanical loading conditions.

Together, the mixed myocardial and systemic effects of SGLTi and the inconsistent observations of SGLTi on contractility underscore the need for a rigorous approach to study human heart tissue contractility under carefully defined preloads/afterloads, and in response to drugs. In this study, we optimized a hLMS platform to study the biomechanical effects of fuels and drugs under a range of physiologic loading conditions to interrogate the hypothesis that SGLT inhibition directly and acutely alters contractility in HCM hLMS distinct from the effects of ketone supplementation.

## METHODS

Full methods are available in the Supplement. Briefly, human myocardial tissue was procured under the protocols and ethical regulations approved by the Institutional Review Boards at the University of Pennsylvania and the Gift-of-Life Donor Program (Philadelphia, PA). Non-failing or HCM tissue was dissected in ice-cold cardioplegia, then 300 µm living myocardial slices were generated in oxygen bubbled Slicing Solution, adhered to custom anchors, and allowed to equilibrate for 1 hour at 37°C in Recording Solution with or without supplemental ketones (see Supplemental Methods for solution components).

Slices were then loaded onto the IonOptix Cardiac Slice System and following a short equilibration period work loops were generated at 4% strain and afterloads of 25%, 50%, and 75% of developed isometric force. Measurements were repeated at 8%, 12%, and 16% strain. Slices were then de-stretched to 0% strain and drug allowed to wash in for 20 minutes. After drug treatment, the same mechanical protocol was performed. Drugs used in this study include: 5 μM isoproterenol (Sigma), 1 μM mavacamten (Medchem Express), 10 μM sotagliflozin (Lexicon Pharmaceuticals) or 10 μM empagliflozin (Cayman Chemical Company).

Raw work loop data were analyzed using IonWizard (IonOptix). Mixed effect linear regression modeling was used to examine associations between clinical parameters, experimental conditions, and contractility parameters. Model outputs are summarized by two parameters: an average at centered strain (i.e. the modeled work or developed stress at 10% strain), and the interaction between the contractile parameter of interest and strain (i.e. change in work per unit increase in strain; work-strain slope or the end diastolic stress strain slope).

## RESULTS

### Clinical Characteristics

We generated hLMS from 32 hearts (16 non-failing donors and 16 septal myectomy specimens from individuals with HCM). 4-8 slices were studied from each heart, totaling 120 slices. Demographic and clinical data are summarized in **Table 1** and in the online data repository. Demographic and clinical characteristics were similar between groups. As expected, individuals from whom HCM samples were obtained demonstrated greater LV wall thickness, higher ejection fraction, and higher use of beta blockers and calcium channel blockers.

**Table 1.** Demographic and Clinical Characteristics of Cardiac Samples Used for hLMS Generation. The location of LV maximal wall thickness was the septum for patients with HCM. p-values obtained from 2-sided t-test. Abbreviations: LVEF: left ventricular ejection fraction. LVEDD: left ventricular end diastolic dimension. ACEi or ARB: angiotensin converting enzyme inhibitor or angiotensin receptor blocker. LVOT: left ventricular outflow tract.

| Variable | Non-Failing (N=16) | HCM Myectomy (N=16) | p-values |
| --- | --- | --- | --- |
| **Demographics** |  |  |  |
| Age, mean (SD) | 54.75 (10.80) | 57.00 (16.30) | 0.64 |
| Male Sex, N (%) | 6 (38) | 9 (56) | 0.31 |
| Race/Ethnicity |  |  |  |
| White, N (%) | 13 (81) | 14 (88) | 0.63 |
| Black or African American, N (%) | 3 (19) | 2 (12) | 0.63 |
| Hispanic, N (%) | 1 (6.25) | 0 (0) | 1.00 |
| **Clinical and Echocardiographic Characteristics** |  |  |  |
| Obesity, N (%) | 8 (50) | 7 (43.75) | 0.73 |
| BMI, mean (SD) | 31.76 (8.25) | 31.94 (8.37) | 0.95 |
| Diabetes, N (%) | 10 (62.5) | 1 (6.25) | **0.001** |
| Hypertension, N (%) | 11 (68.75) | 10 (62.5) | 0.72 |
| LVEF, mean % (SD) | 60.44 (9.30) | 65.31 (5.95) | 0.09 |
| LV Mass Index, mean g/m<sup>2</sup> (SD) | 111.03 (19.72) | 134.59 (38.55) | **0.03** |
| LV Maximal Wall Thickness, mean cm (SD) | 0.98 (0.22) | 2.15 (0.43) | **<0.0001** |
| LVEDD, mean cm (SD) | 4.53 (0.87) | 4.24 (0.61) | 0.28 |
| **Medications** |  |  |  |
| Beta Blocker, N (%) | 3 (18.75) | 13 (81.25) | **0.0006** |
| ACEi or ARB, N (%) | 4 (64) | 6 (37.5) | 0.46 |
| Calcium Channel Blocker, N (%) | 3 (18.75) | 5 (31.25) | 0.69 |
| Diuretic, N (%) | 1 (6.25) | 2 (12.5) | 1.00 |
| SGLTi, N (%) | 0 (0) | 0 (0) |  |
| **HCM Specific Characteristics** |  |  |  |
| LVOT Gradient |  |  |  |
| Rest, mean mmHg (SD) | N/A | 69.75 (40.17) |  |
| Valsalva, mean mmHg (SD) | N/A | 96.53 (45.21) |  |
| Genetic Testing |  |  |  |
| No, N (%) | N/A | 8 (50) |  |
| Yes, N (%) | N/A | 8 (50) |  |
| Positive, N (%) |  | 1 (6) |  |
| Negative, N (%) |  | 6 (38) |  |
| Inconclusive, N (%) |  | 1 (6) |  |

### Influence of Clinical Variables on Myocardial Work

We assessed baseline work across predetermined load conditions of hLMS from both non-failing donors (NF) and septal myectomy tissue from individuals with HCM. (**Figure 1A**). Principal components analysis revealed human level slice-to-slice variability with separation between NF vs. HCM slices. Examination of the raw baseline work loops showed that across all load conditions, but particularly at low preloads, average work loops and quantified work from HCM slices were larger than from NF (**Figure 1D-E, S2-S3**).

**Figure 1.**
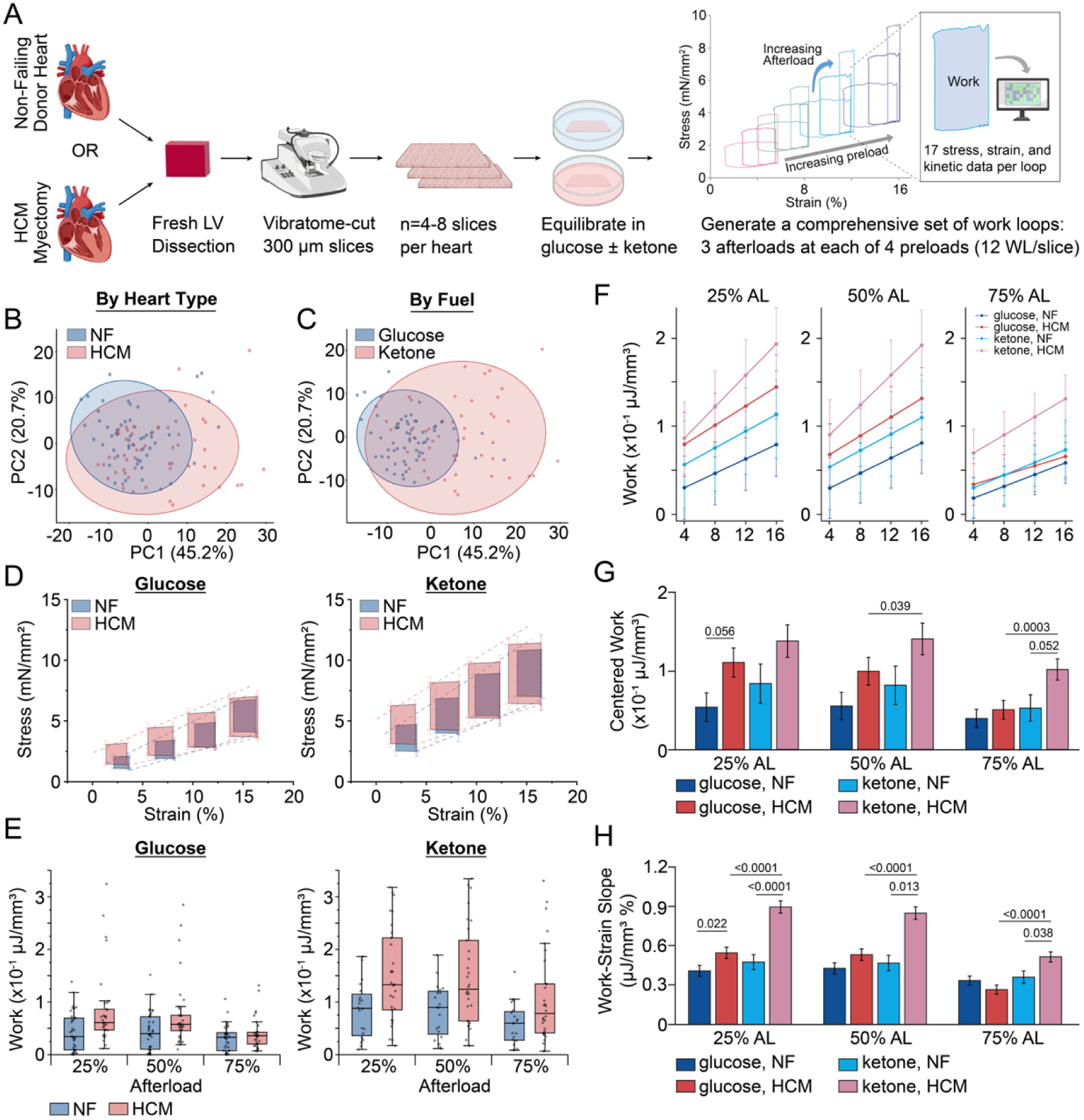
Baseline Work Assessment of hLMS from Non-Failing or HCM Tissue in Glucose or Glucose Supplemented with Ketones. **A.** Workflow for generation and analysis of hLMS. Freshly explanted human heart tissue from either non-failing donors or from HCM individuals following surgical myectomy are brough to the lab and slices are immediately generated. hLMS are equilibrated in either glucose only containing buffer or glucose and ketone containing buffer. A set of work loops at prespecified preloads and afterloads are generated and processed for analysis. Principal components analysis of all work loop parameter data segmented either by **B.** heart type or **C.** fuel. Each dot represents one slice and shaded circles represent 95% confidence ellipses. **D**. Average work loops ± standard error from the mean (SEM) at all preloads and 50% afterload. Dotted lines represent the end diastolic and end systolic stress strain relationships generated from the full set of 12 work loops. **E**. Calculated work at 8% strain for each slice at 25, 50, and 75% afterloads. Boxes denote median, 25^th^ and 75^th^ percentile and whiskers represent 1.5 interquartile range (IQR). **F.** Mixed effect linear model output for work across all preloads and afterloads (Mean ± standard error). **G.** Centered work (modeled work at 10% strain) across all baseline conditions at 25, 50, and 75% afterloads. Values denote significant p-values (<0.05) from mixed-effect linear regression and *pwcompare* for comparison between two groups at centered strain. **H.** Work-strain slope across all baseline conditions at 25, 50, and 75% afterloads. Values denote significant p-values (<0.05) for mixed-effect linear regression and *pwcompare* comparisons between two groups across strains.

We next assessed the relationship between clinical characteristics and work generated by hLMS with our mixed effect linear regression model (**Figure 1F**). For NF slices across afterloads, older age was inversely associated with both work at centered strain as well as the capacity for hLMS to increase work with increased strain (work-strain slope). Presence of hypertension was associated with increased work-strain slope in both NF and HCM hLMS. Ejection fraction was positively correlated with work in NF but not HCM hLMS. In HCM hLMS, greater LVMI was associated with increased work-strain slope at low and medium afterloads (**Table 2**).

**Table 2:** Dependence of Work and Work-Strain Slope on Demographic and Clinical Variables. Mixed effects linear model coefficients (β) and p-values of association for centered work (10% strain) and work-strain slope across heart types (NF or HCM) and all three afterloads. p-values obtained from STATA mixed effect linear regression modeling and *pwcompare. p-values* less than 0.05 are bolded.

|  | 25% Afterload |  | 50% Afterload |  | 75% Afterload |  |
| --- | --- | --- | --- | --- | --- | --- |
| Variable | $\beta$ (SEM) | p-value | $\beta$ (SEM) | p-value | $\beta$ (SEM) | p-value |
| Age |  |  |  |  |  |  |
| NF, work | -3.1 (0.9)x10<sup>-3</sup> | **0.0008** | -3.2 (0.9)x10<sup>-3</sup> | **0.0004** | -2.2 (0.6)x10<sup>-3</sup> | **0.0001** |
| HCM, work | 0.8 (1.6)x10<sup>-3</sup> | 0.63 | 0.8 (1.5)x10<sup>-3</sup> | 0.59 | 0.3 (0.6)x10<sup>-3</sup> | 0.65 |
| NF, slope | -2.1 (0.4)x10<sup>-2</sup> | **<0.0001** | -2.0 (0.4)x10<sup>-2</sup> | **<0.0001** | -1.5 (0.4)x10<sup>-2</sup> | **<0.0001** |
| HCM, slope | -0.3 (0.2)x10<sup>-2</sup> | 0.11 | -0.3 (0.2)x10<sup>-2</sup> | 0.11 | -0.1 (0.2)x10<sup>-2</sup> | 0.51 |
| Sex |  |  |  |  |  |  |
| NF, work | -1.5 (2.4)x10<sup>-2</sup> | 0.52 | -2.0 (2.3)x10<sup>-2</sup> | 0.37 | -1.8 (1.4)x10<sup>-2</sup> | 0.21 |
| HCM, work | 3.7 (5.4)x10<sup>-2</sup> | 0.48 | 4.2 (4.9)x10<sup>-2</sup> | 0.39 | 2.0 (2.0)x10<sup>-2</sup> | 0.32 |
| NF, slope | -1.6 (1.0)x10<sup>-1</sup> | 0.12 | -1.7 (1.1)x10<sup>-1</sup> | 0.10 | -2.5 (1.0)x10<sup>-1</sup> | **0.009** |
| HCM, slope | -1.6 (0.7)x10<sup>-1</sup> | **0.021** | -1.2 (0.7)x10<sup>-1</sup> | 0.10 | -0.6 (0.7)x10<sup>-1</sup> | 0.36 |
| BMI |  |  |  |  |  |  |
| NF, work | -1.3 (1.1)x10<sup>-3</sup> | 0.26 | -1.0 (1.1)x10<sup>-3</sup> | 0.34 | -0.4 (0.6)x10<sup>-3</sup> | 0.53 |
| HCM, work | -1.7 (2.6)x10<sup>-3</sup> | 0.52 | -0.8 (2.3)x10<sup>-3</sup> | 0.74 | 0.6 (0.9)x10<sup>-3</sup> | 0.49 |
| NF, slope | -7.7 (4.6)x10<sup>-3</sup> | 0.094 | -6.6 (4.7)x10<sup>-3</sup> | 0.15 | -2.4 (4.3)x10<sup>-3</sup> | 0.57 |
| HCM, slope | -0.1 (2.8)x10<sup>-3</sup> | 0.98 | 1.8 (2.9)x10<sup>-3</sup> | 0.53 | 6.0 (2.8)x10<sup>-3</sup> | **0.034** |
| Diabetes |  |  |  |  |  |  |
| NF, work | -0.2 (2.1)x10<sup>-2</sup> | 0.92 | -0.1 (2.0)x10<sup>-2</sup> | 0.95 | 0.2 (1.3)x10<sup>-2</sup> | 0.89 |
| HCM, work | -3.1 (11)x10<sup>-2</sup> | 0.78 | -1.6 (10)x10<sup>-2</sup> | 0.87 | 2.7 (4.2)x10<sup>-2</sup> | 0.52 |
| NF, slope | 1.4 (0.9)x10<sup>-1</sup> | 0.11 | 1.8 (0.9)x10<sup>-1</sup> | **0.044** | 1.4 (0.8)x10<sup>-1</sup> | 0.078 |
| HCM, slope | -2.6 (1.4)x10<sup>-1</sup> | 0.068 | -2.5 (1.4)x10<sup>-1</sup> | 0.081 | -0.6 (1.4)x10<sup>-1</sup> | 0.62 |
| Hypertension |  |  |  |  |  |  |
| NF, work | 5.9 (2.7)x10<sup>-2</sup> | **0.031** | 6.1 (2.6)x10<sup>-2</sup> | **0.017** | 4.1 (1.6)x10<sup>-2</sup> | **0.010** |
| HCM, work | 2.5 (5.4)x10<sup>-2</sup> | 0.64 | 2.2 (4.8)x10<sup>-2</sup> | 0.64 | 0.0 (2.0)x10<sup>-2</sup> | 0.99 |
| NF, slope | 2.9 (1.1)x10<sup>-1</sup> | **0.010** | 2.8 (1.1)x10<sup>-1</sup> | **0.016** | 3.1 (1.1)x10<sup>-1</sup> | **0.004** |
| HCM, slope | 3.2 (0.7)x10<sup>-1</sup> | **<0.0001** | 3.3 (0.7)x10<sup>-1</sup> | **<0.0001** | 1.9 (0.7) x10<sup>-1</sup> | **0.016** |
| LVEF |  |  |  |  |  |  |
| NF, work | 1.7 (0.9)x10<sup>-3</sup> | **0.039** | 1.6 (0.8)x10<sup>-3</sup> | **0.047** | 0.9 (0.5)x10<sup>-3</sup> | 0.067 |
| HCM, work | -3.9 (5.9)x10<sup>-3</sup> | 0.50 | -4.0 (5.3)x10<sup>-3</sup> | 0.45 | -3.3 (2.3)x10<sup>-3</sup> | 0.14 |
| NF, slope | 0.5 (0.4)x10<sup>-2</sup> | 0.13 | 0.6 (0.4)x10<sup>-2</sup> | 0.12 | 0.6 (0.3)x10<sup>-2</sup> | 0.072 |
| HCM, slope | 1.2 (0.7)x10<sup>-2</sup> | 0.10 | 1.1 (0.8)x10<sup>-2</sup> | 0.11 | 0.4 (0.7)x10<sup>-2</sup> | 0.61 |
| LVMI |  |  |  |  |  |  |
| NF, work | 9.7 (6.0)x10<sup>-4</sup> | 0.10 | 9.1 (5.7)x10<sup>-4</sup> | 0.11 | 5.3 (3.6)x10<sup>-4</sup> | 0.14 |
| HCM, work | -6.3 (8.1)x10<sup>-4</sup> | 0.44 | -5.5 (7.4)x10<sup>-4</sup> | 0.45 | 0.0 (3.0)x10<sup>-4</sup> | 0.99 |
| NF, slope | 0.8 (2.6)x10<sup>-3</sup> | 0.74 | 0.5 (2.6)x10<sup>-3</sup> | 0.85 | 2.2 (2.4)x10<sup>-3</sup> | 0.36 |
| HCM, slope | 4.3 (0.9)x10<sup>-3</sup> | **<0.0001** | 3.2 (0.9)x10<sup>-3</sup> | **0.0012** | 1.5 (0.9)x10<sup>-3</sup> | 0.27 |

Quantification of centered work reveals overall increased work and a higher work-strain slope in HCM slices relative to NF at low afterloads (**Figure 1G,H**). Developed stress, the total amount of normalized force generated during isometric systolic contraction, was significantly higher in HCM relative to NF slices at all afterloads (**Table S2**). Taken together, these data confirm increased contractility in HCM relative to NF.

### Ketone Supplementation Augments Contractile Work

We next assessed the impact of ketone supplementation by comparing slices in either glucose-only Recording Solution or a Recording Solution supplemented with 6mM 3-OHB (Ketone). Ketone supplementation augmented contractility particularly in HCM slices with greatest effect at higher afterloads, and increased work-strain slope across all load conditions (**Figure 1G,H**).

Principle component analysis showed clear separation between glucose and ketone fueled slices (**Figure 1B**). Ketone supplementation also substantially augmented the size of average work loops regardless of underlying disease etiology (**Figure 1D, S2**). Quantified work across load conditions demonstrated that ketone supplementation increases contractility in both NF and HCM. (**Figure 1E, S3**).

Quantification of mixed effects linear regression model outputs revealed that work at centered strain in ketone supplemented media was significantly higher than glucose alone in HCM slices at moderate (50%) and high (75%) afterloads, but not at the lowest afterload (**Figure 1G**). The work-strain slope in HCM slices was significantly higher in ketone than in glucose alone at all afterloads (**Figure 1H**). We further observe that when ketones are supplemented, HCM slices relative to NF slices have greater centered work at high afterloads and greater work-strain slope at all afterloads. Together, these data reveal the positive inotropic effect of ketones particularly in the context of HCM.

We performed targeted metabolomics in a subset of slices to confirm the metabolic impact of ketone supplementation. As expected, beta-hydroxybutyrate was significantly increased in ketone-supplemented slices. We further observed an increase in C4-OH Butyryl in ketone supplemented slices suggestive of increased ketone metabolism. There was a compensatory trend towards reduced medium and long chain acylcarnitines (**Table S2**).

### Validation of hLMS Drug Testing Protocol

Recognizing intrinsic slice-to-slice variability, we developed a methodology to conduct paired analyses of hLMS before and after acute addition of a drug (**Figure 2A**). To validate the platform, hLMS were recirculated either with dimethylsulfoxide (DMSO) serving as an inert control, isoproterenol (5 μM), a known positive inotrope, or mavacamten (1 μM), a myosin modulator that reduces contractility. PCA analysis on fold change in work loop parameters following drug administration relative to baseline shows a separation between DMSO, isoproterenol, and mavacamten treated hLMS (**Figure 2B**).

**Figure 2.**
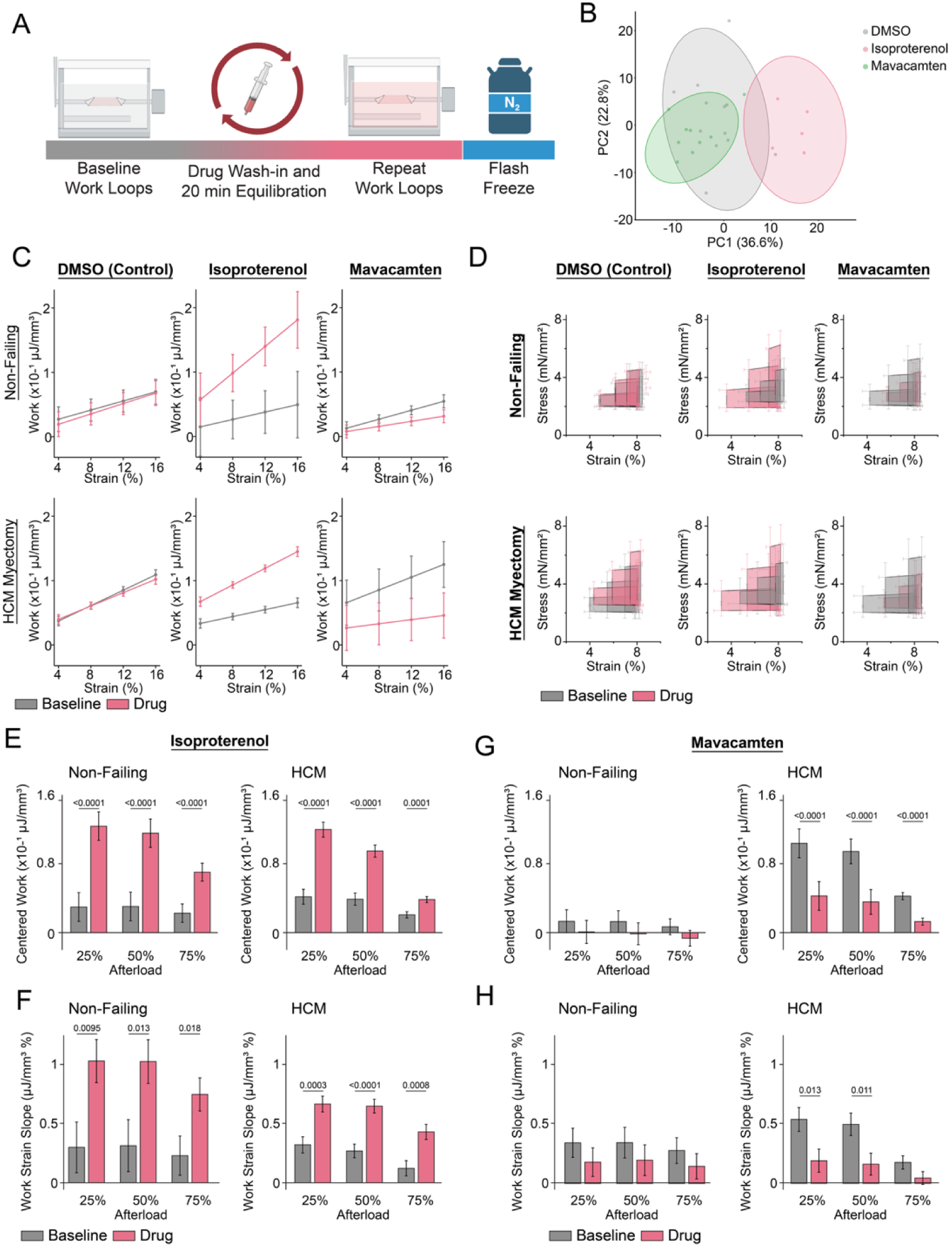
Validation of Human Living Myocardial Slice Work Loops as a Platform to Test Contractility-Modulating Drugs. **A.** Workflow for drug testing. Following baseline measurement of baseline work loops, LMS is destretched and drug is washed in for 20 minutes. After wash-in, work loop measurements are repeated in the presence of drug. **B.** PCA of the fold change of work loop parameter data between the drug run and baseline run of work loops. Each dot represents one slice and shaded circles represent 95% confidence ellipses. **C.** Mixed effect linear model output for work across all preloads at 50% afterload (Mean ± standard error) at baseline (gray) and following drug (pink). **D.** Average work loops ± standard error from the mean (SEM) at all afterloads and 8% strain. **E.** Centered work and **F.** Work strain slope across all loads and heart types at baseline and in response to isoproterenol. **G.** Centered work and H. Work strain slope across all loads and heart types at baseline and in response to mavacamten. For E-G, bars represent mean ± standard error and values denote p-values between two groups using mixed effects linar regression modeling, *pwcompare,* and *lincom* to generate custom contrasts of regression slopes across conditions.

Mixed effects linear regression modeling of work revealed an overall null effect of DMSO, increased work with isoproterenol, and decreased work with mavacamten (**Figure 2C, S4-S6**). To determine if these were reflected in the raw data, we examined average work loops as well as raw work of drug-treated hLMS relative to baseline (**Figure 2D, S7-S10**). We again confirmed that baseline work loops overlapped well with DMSO-treated loops, suggesting minimal effect of repeated measurements on work. Isoproterenol work loops were larger and mavacamten smaller than their respective baselines, consistent with their known drug effects.

We next examined the effects of DMSO, isoproterenol, or mavacamten in modeled centered work, work-strain slope, or centered active developed stress in NF or HCM across all afterloads. DMSO treatment did not change work at centered strain or work-strain slopes regardless of afterload or tissue source (NF or HCM) (**Figure S11**). There was slight but significant increase in developed stress following DMSO treatment (**Table S4**). Conversely, isoproterenol-treated hLMS had increased work at centered strain, work-strain slope, and developed stress regardless of afterload or underlying disease, confirming its positive inotropic effect (**Figure 2E**). The magnitude of increase was blunted in HCM hLMS compared to NF. Statistical comparison between differences in work at centered strain and work-strain slope of isoproterenol- vs DMSO-treated hLMS further confirms significant positive inotropy of isoproterenol (**Table S5**).

Following mavacamten treatment, NF hLMS showed a trend towards a reduction in work and a modest reduction in developed stress. Conversely, mavacamten treated HCM hLMS displayed significantly reduced work and developed stress regardless of load, suggesting a disease-specific drug effect. The magnitude of drug effect on HCM hLMS was greatest at low afterloads but absolute work was lowest at high afterloads. When statistically comparing mavacamten treatment to DMSO, we observed a significant effect on work but not work-strain slope in HCM hLMS (**Table S4**), suggesting that mavacamten treatment does not significantly affect length-dependent activation of myocardium.

These data therefore confirm the ability to repeat measurements on hLMS following drug treatment, recapitulate key positive and negative contractile effects of drugs, and identify both disease and load-associated drug effects.

### SGLT inhibition acutely and directly reduces contractile work

We next explored the effects of two SGLT inhibitors on contractility. We studied sotagliflozin (10 μM), a dual SGLT1 and SGLT2 inhibitor, and empagliflozin (10 μM), a selective SGLT2 inhibitor. Principal components analysis of slices treated with DMSO or either sotagliflozin or empagliflozin relative to their baseline displayed significant overlap, suggesting overall modest effects of SGLT inhibition (**Figure S12**). Average work loops were smaller following SGLTi treatment relative to baseline in both NF and HCM hLMS (**Figure 3A,B, S13**). Paired baseline to treated plots of raw work data showed a consistent reduction in work in most hLMS (**Figure 3C,D, S14-15)**.

**Figure 3.**
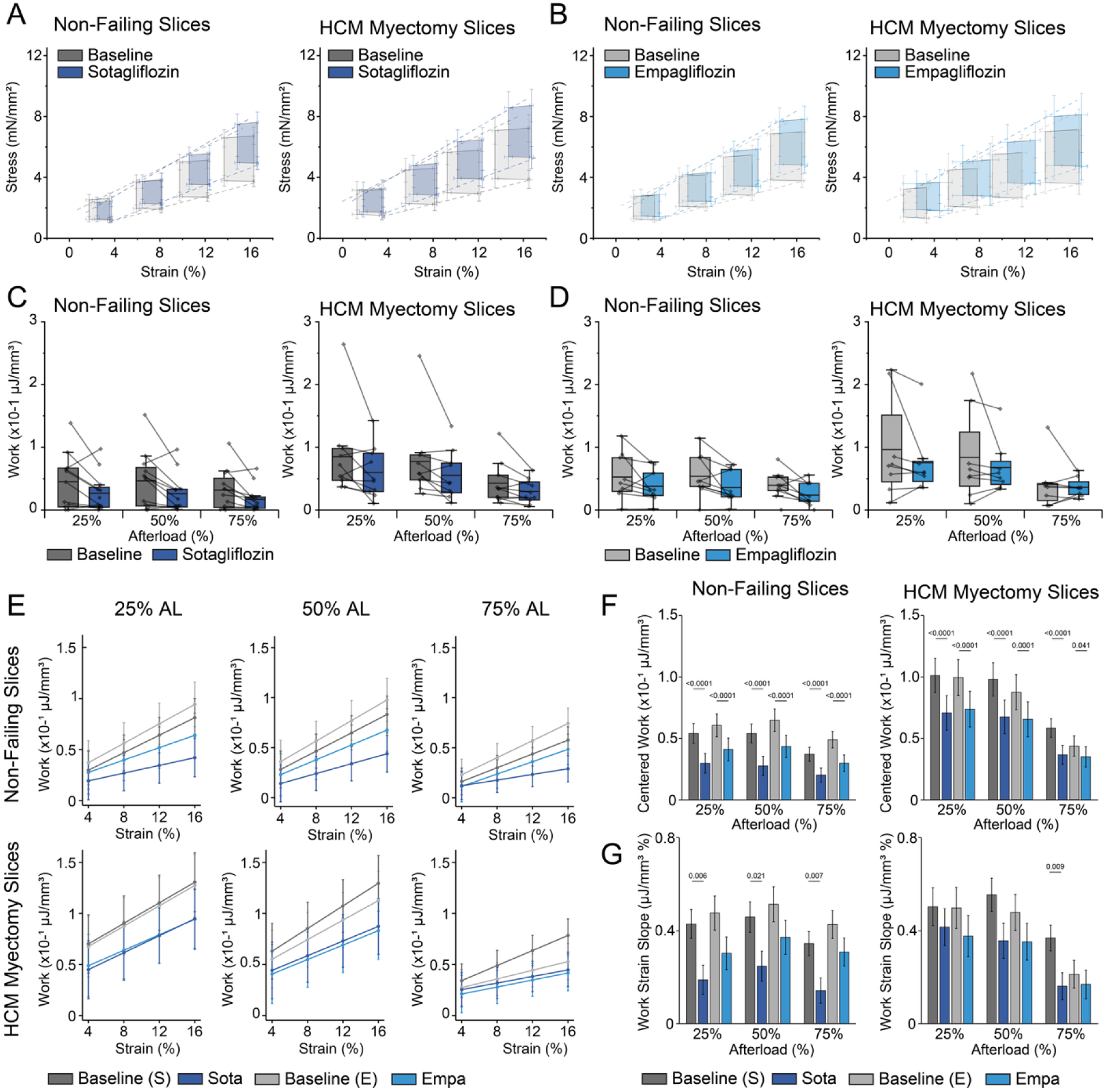
Acute and Direct Effects of SGLTi on Work in Human Living Myocardial Slices. Average work loops ± standard error from the mean (SEM) at all preloads and 50% afterload for **A.** slices treated with sotagliflozin and **B.** slices treated with empagliflozin. Dotted lines refer to the end systolic and end diastolic stress strain relationships. **C-D.** Paired work measurements at baseline and **C.** in response to sotagliflozin or **D**. in response to empagliflozin. Boxes denote median, 25^th^ and 75^th^ percentile and whiskers represent 1.5 interquartile range (IQR). **E**. Mixed effect linear model output at baseline (S refers to eventual sotagliflozin treated slices and E refers to eventual empagliflozin treated slices) and following treatment with SGLTi. Displayed as mean ± standard error. **F.** Centered work ± standard error and **G.** work strain slope ± standard error at all afterloads. Values refer to p-values comparing drug to baseline using mixed effects linear regression modeling, *pwcompare,* and *lincom* to generate custom contrasts of regression slopes across conditions.

Linear modeling confirmed modest but significant reductions in centered work and developed stress across all afterloads (**Figure 3E,F, S16, Table S6**) conferred by SGLTi. The magnitude of reduction in work was similar between NF and HCM hLMS. In NF slices, sotagliflozin significantly reduced work-strain slope across all afterloads. Likewise, in HCM hLMS, sotagliflozin significantly reduced work-strain slope at high afterloads. Empagliflozin-treated slices behaved similarly, however reductions in work-strain slope were not statistically significant (**Figure 3G**). When statistically comparing SGLTi treatment with DMSO, there were significant differences between either sotagliflozin or empagliflozin and DMSO (**Figure S17**). At high afterloads, only sotagliflozin was significantly different from DMSO, and the effect of sotagliflozin was statistically significantly different from empagliflozin. (**Figure S17**). Together these data show a largely concordant reduction of work regardless of which SGLTi was administered, with only sotagliflozin reducing work in HCM hLMS at high afterloads.

We further explored the effect of SGLTi on diastolic function. Linear regression modeling of isovolumic relaxation time (IVRT) demonstrated overall minimal changes when treated with SGLTi. There was significantly faster IVRT in HCM slices treated with sotagliflozin at medium afterload (**Table 3**). We also assessed the effect of SGLTi on end-diastolic stress-strain relationship (EDSSR). To do this, we performed mixed effects linear regression modeling on end-diastolic stress at prespecified strains. With the interaction term between end diastolic stress and strain, we assessed the slope of the end-diastolic stress strain regression. This slope is comparable to the end-diastolic pressure volume relationship (EDPVR), a key metric of diastolic function. We observed a significant increase in the end-diastolic stress strain slope with both DMSO treatment (our null control) as well as either SGLTi. However, when comparing across drugs, there were no differences between DMSO treatment and either SGLTi, suggesting no significant effect of SGLTi on diastolic function in this model (**Table S7, Figure S18**).

**Table 3:**
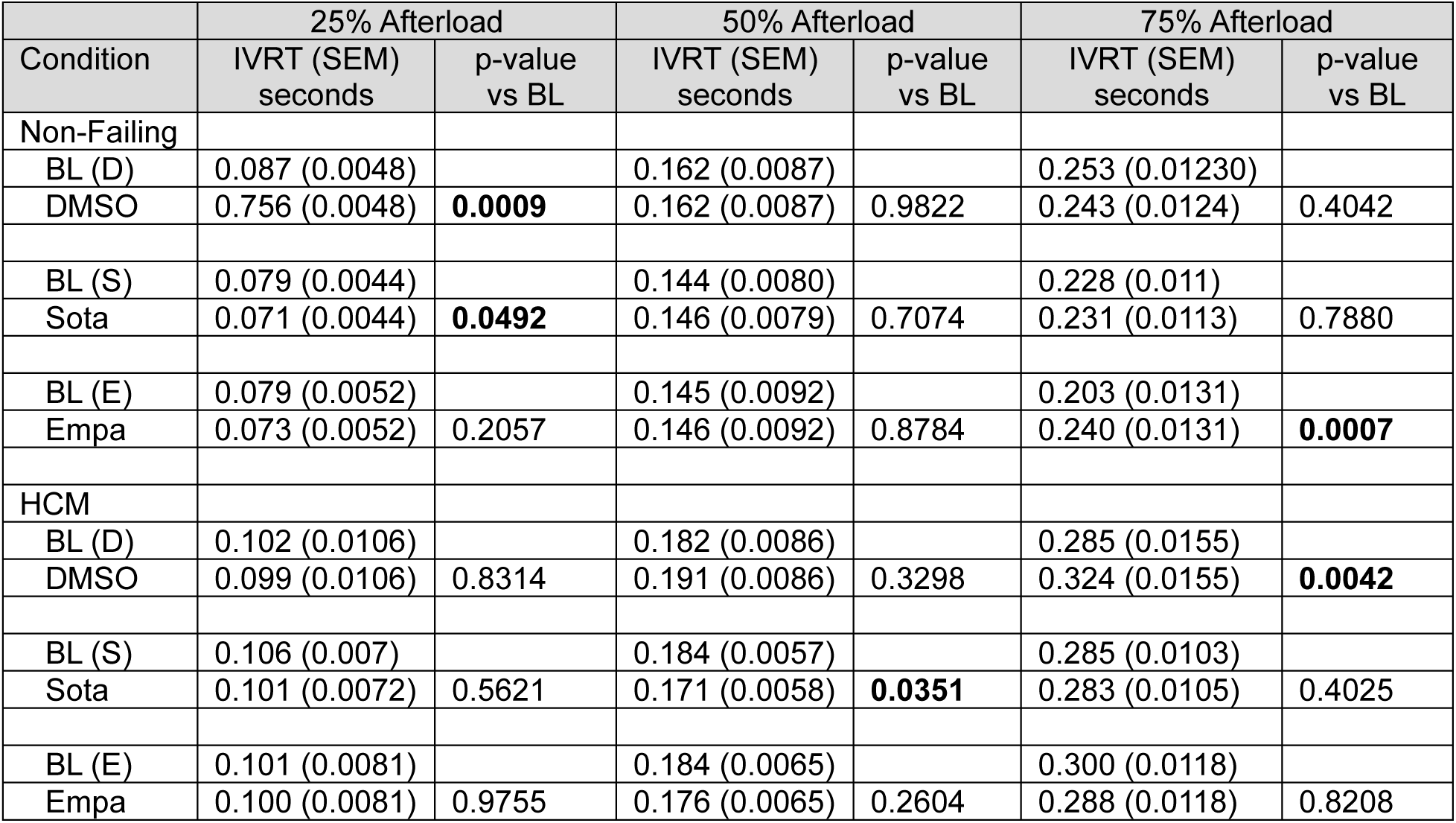
Mixed Linear Regression Modeling of Isovolumic Relaxation Time. . Values are denoted as mean (standard error). p-values show mixed effects linear regression modeling comparison between DMSO or SGLTi treated run with baseline. p values less than 0.05 are bolded.

|  | 25% Afterload |  | 50% Afterload |  | 75% Afterload |  |
| --- | --- | --- | --- | --- | --- | --- |
| Condition | IVRT (SEM)<br>seconds | p-value<br>vs BL | IVRT (SEM)<br>seconds | p-value<br>vs BL | IVRT (SEM)<br>seconds | p-value<br>vs BL |
| Non-Failing |  |  |  |  |  |  |
| BL (D) | 0.087 (0.0048) |  | 0.162 (0.0087) |  | 0.253 (0.01230) |  |
| DMSO | 0.756 (0.0048) | **0.0009** | 0.162 (0.0087) | 0.9822 | 0.243 (0.0124) | 0.4042 |
| BL (S) | 0.079 (0.0044) |  | 0.144 (0.0080) |  | 0.228 (0.011) |  |
| Sota | 0.071 (0.0044) | **0.0492** | 0.146 (0.0079) | 0.7074 | 0.231 (0.0113) | 0.7880 |
| BL (E) | 0.079 (0.0052) |  | 0.145 (0.0092) |  | 0.203 (0.0131) |  |
| Empa | 0.073 (0.0052) | 0.2057 | 0.146 (0.0092) | 0.8784 | 0.240 (0.0131) | **0.0007** |
| HCM |  |  |  |  |  |  |
| BL (D) | 0.102 (0.0106) |  | 0.182 (0.0086) |  | 0.285 (0.0155) |  |
| DMSO | 0.099 (0.0106) | 0.8314 | 0.191 (0.0086) | 0.3298 | 0.324 (0.0155) | **0.0042** |
| BL (S) | 0.106 (0.007) |  | 0.184 (0.0057) |  | 0.285 (0.0103) |  |
| Sota | 0.101 (0.0072) | 0.5621 | 0.171 (0.0058) | **0.0351** | 0.283 (0.0105) | 0.4025 |
| BL (E) | 0.101 (0.0081) |  | 0.184 (0.0065) |  | 0.300 (0.0118) |  |
| Empa | 0.100 (0.0081) | 0.9755 | 0.176 (0.0065) | 0.2604 | 0.288 (0.0118) | 0.8208 |

To contextualize our observations on contractility, we performed targeted metabolomics on a subset of hLMS treated with SGLTi. Metabolomics analysis of hLMS treated with sotagliflozin or empagliflozin compared to DMSO revealed no significant changes on the abundance of any metabolites (**Table S8**). These results suggest that reduced contractility following SGLTi treatment occurs *without* significantly changing the metabolic landscape.

## DISCUSSION

This study addresses a longstanding need for *ex vivo* multicellular models of human cardiac contractility that reflect patient-level data and allow pharmacologic testing under well-defined loading conditions.^38,39^ We developed a methodology for phenotyping human living myocardial slices under multiple loads, fuels, and drug exposures, then leveraged the system to reveal the distinct effects of ketone supplementation and SGLTi in HCM.

### hLMS provide a novel platform to study load-dependent HCM physiology

While prior studies demonstrate load-dependent effects on contractility and gene expression in cultured rat LMS,^14,40^ prior work in human slices has relied on isometric or auxotonic culture and predominantly focused on feasibility of long-term maintenance.^16,17,38^ The capacity to interrogate load-dependent physiology is an important attribute of this human model system. In a recent systematic review of 602 HCM studies across species, only a minority of models recapitulate multiple hallmarks of HCM, and none recapitulate afterload elevation.^41^ Since short-term culture of LMS significantly remodels metabolic function, contractility, and gene expression,^18,42^ we studied hLMS acutely after explantation. We observe hypercontractility in HCM LMS, a central pathophysiologic feature of HCM, thought to be an adaptive response to increased afterload from left ventricular outflow tract obstruction or increased wall stress in the absence of obstruction. This is known as the Anrep effect where afterload-dependent increased recruitment of myosin heads is compounded by destabilization of myosin’s resting configuration.^43,44^ Interestingly, hypercontractility is blunted at high afterloads in glucose-only HCM LMS, potentially reflecting reserve depletion where prolonged exposure to high afterload drains myocardial energy reserves.^44^ Ketone supplementation restores hypercontractility of HCM slices at high afterload, potentially demonstrating increased reliance of HCM tissue on alternate energy sources. While the current study focuses on HCM pathophysiology, our model’s capacity to study load-dependence is broadly applicable for other disease states with afterload mismatch, including aortic stenosis or hypertensive cardiomyopathy.

Using mixed linear regression modeling, we probed associations between hLMS myocardial work and clinical characteristics of individuals from whose tissue hLMS were generated. In NF hLMS, increasing age was inversely associated with work and the capacity to augment work with strain. This is consistent with clinical decline in cardiac reserve seen with increasing age,^45^ and supports use of hLMS for further *ex vivo* studies of aging. Notably, this age-associated decreased contractility is not seen in HCM, consistent with the notion that genetically driven hypercontractility overrides the contribution of age. Presence of hypertension predicts hLMS ability to augment contractility with increasing strain, consistent with compensatory remodeling associated with increased afterload. Together, these data strongly support use of fresh hLMS as a model of *in vivo* contractility that reflects antecedent clinical data.

### Ketone supplementation increases hLMS contractility

Ketone supplementation had a positive inotropic effect on hLMS across all afterloads. While ketone supplementation has previously been shown to acutely increase cardiac output *in vivo*, this occurred in the context of reduced systemic vascular resistance.^30,46^ We previously found that ketone supplementation increases contractility in isolated, unloaded human cardiomyocytes.^47^ With the capacity to modify load in the hLMS, we were able to ascertain that ketones acutely and directly increase myocardial contractility. The most marked inotropic effects of ketones occurred in HCM hLMS, consistent with increased myocardial demand and reduced metabolic flexibility in HCM leading to greater reliance on ketone metabolism.^48^ Increased levels of C4-OH Butyryl confirms increased ketone metabolism in ketone-supplemented slices. The trend towards reduced medium and long chain acylcarnitines is potentially indicative of substrate competition and altered fatty-acid flux into mitochondria. Future studies may use respirometry-based approaches to assess altered substrate handling to better define the metabolic effects of ketones in HCM and NF tissues.

### Disease- and load-dependent responses to positive and negative inotropes

Beyond demonstrating known positive inotropy of isoproterenol in hLMS to validate the model, we observed a blunted effect in HCM hLMS particularly at high afterloads, consistent with impaired contractile reserve and adrenergic responsiveness in HCM. Disease-associated drug effects were more striking with mavacamten, where we observed a much more substantial reduction in contractility in HCM hLMS compared to NF. This is consistent with the notion that the effect of mavacamten, which stabilizes the myosin super-relaxed state (SRX), is dependent on the ratio of disordered relaxed state (DRX) myosin to SRX which is increased in HCM compared to normal hearts.^49–51^ We further observed that mavacamten had the greatest magnitude of effect to reduce contractility at low afterloads in HCM hLMS, consistent with the biophysical properties of mechanical load affecting the dynamics of the myosin motor.^52,53^ Together, these data validate use of hLMS as a drug testing platform capable of elucidating disease- and load-dependent effects on contractility.

### SGLTi reduce work in hLMS

We found a modest reduction of work following acute administration of SGLTi, consistent with prior studies using HCM engineered heart tissues (EHTs).^37^ Given the pathologic hypercontractility of HCM tissue, this would be expected to be a favorable therapeutic effect. Interestingly, our results contrast with prior work in both human trabeculae^54^ and living myocardial slices^36^ predominantly from failing hearts, which revealed no effect on active contractility from SGLTi, as well as in isolated cardiomyocytes, which revealed increased fractional shortening from SGLTi.^34^ It is possible that SGLTi have different effects depending on load and the contractile state at baseline.

Treatment of the HCM hLMS with sotagliflozin, but not empagliflozin, reduced IVRT at medium afterload, potentially indicating more efficient relaxation kinetics under certain loading conditions. These results are consistent with results from Wjinker et al that demonstrate decreased relaxation time in HCM EHTs treated with SGLTi.^37^ We observe no effect on the end diastolic stress-strain slope (i.e. diastolic stiffness) across a range of physiological afterloads when compared to DMSO control. However, this result is confounded by the observation that diastolic stiffness *increased* in all slices regardless of treatment, potentially due to time in a less stretched state during drug loading leading to contraction of the slice.^17^ Our diastology results again differ from studies done in trabeculae from failing hearts.^54^ The effects on passive stiffness in trabeculae were clearest at maximally stretched, supraphysiologic sarcomere lengths, whereas we restricted preload to physiologic ranges.

One potential mechanism for decreased contractility in HCM slices following SGLTi treatment is altered NCX activity. In healthy hearts, NCX maintains calcium homeostasis by removing calcium from the cell (“forward mode”) with each beat. NCX is thought to have increased “reverse” mode in HCM, leading to increased cytosolic calcium.^37,55^ Wijnker et al demonstrated reduced cytosolic [Ca^2+^] in HCM EHT but not control EHTs treated with SGLTi. While they did not identify a mechanism, their data indicate that altered NCX activity likely underlies acutely decreased contractility and reduced relaxation time seen in HCM EHTs treated with SGLTi. Their data supported the hypothesis that SGLTi may act to block reverse mode and/or increase forward mode NCX activity in HCM.^37^

Clinical translation of disease-modifying therapies for HCM remains low, particularly for non-obstructive HCM. Many therapies have failed to show benefit, including Mavacamten in the recently published ODYSSEY trial, where >20% of patients dropped their ejection fraction below the desired level resulting in drug interruption.^7^ Treatment with SGLTi may confer a more modest effect on reducing myocardial work and lowering the energy requirement. The SONATA trial is currently enrolling patients with HCM with both obstructive and non-obstructive physiology to examine safety and efficacy of sotagliflozin in HCM. The results of our study support the continued assessment of SGLTi in clinical trials in patients with HCM.^56^

### Sotagliflozin demonstrates increased potency compared to empagliflozin in hLMS

Similar to other studies using multiple SGLTi agents,^37,54^ we found largely concordant effects between sotagliflozin and empagliflozin with some important distinctions. In HCM hLMS at high afterloads, we observed a more marked effect of sotagliflozin compared to empagliflozin. The increased potency of sotagliflozin may be explained by sotagliflozin’s greater hydrophobicity compared to empagliflozin, leading to higher membrane affinity^57^ and potentially increasing its overall proximity and activity near reported targets such as sodium channels.^33,37^ This may be relevant in the context of HCM where wall stresses are elevated with or without the additional afterload from left ventricular outflow tract obstruction.

While SGLT2 is not present in human myocardium, previous studies report that SGLT1 is present in human heart tissues.^58^ This is one potential explanation for the increased potency of sotagliflozin, a dual SGLT1/2 inhibitor, in human heart tissues, as compared to empagliflozin, which is more selective for SGLT2. However, while SGLT1 transcripts (*SLC5A1*) have been detected in the Myocardial Applied Genomics Network (MAGNet) repository, we were unable to detect SGLT1 protein in human heart tissue by either western blot or mass spectrometry (data not shown). While there may be low levels of SGLT1 protein in human heart tissue, we believe it is more likely that SGLTi, including sotagliflozin, impart their direct effects on the myocardium through off-target mechanisms, as has been previously shown by other groups.^34^

### Ketone supplementation and SGLTi treatment have distinct and discordant effects

Reduced work following SGLTi administration occurs without affecting ketone levels, confirming that SGLTi and ketones have distinct effects on myocardial contractility. Given the modest ketosis observed with SGLTi administration *in vivo*, one open hypothesis is that the cardioprotective mechanism of SGLTi is due to induced ketosis.^59^ Our data clearly show that the direct impact of SGLTi on contractile function is distinct and discordant from increasing ketones. Moreover, recent evidence in mice that ketosis and SGLTi have different metabolic effects^60^ further supports that SGLTi and ketone supplementation have distinct and opposing mechanisms on contractile function.^61^

### Limitations

Our study has several important limitations. We were underpowered to assess the effect of HCM genotype on contractility parameters as only one patient had a known sarcomere variant. This is not unexpected given the lower rate of obstructive physiology in patients with pathogenic sarcomere variants.^62^ There were high rates of comorbidities including diabetes in the non-failing group, potentially confounding the assessment of SGLT2i in this population. While our use of human tissue is an important advantage over mouse or human induced pluripotent stem cell-derived cardiomyocytes that often fail to recapitulate multiple hallmarks of HCM,^41^ human tissue data is inherently more heterogeneous, as reflected by the high variability in our baseline data. Our use of paired measurements and statistical modeling with random intercepts for heart and slice mitigates some of this variability. Further, given the limited availability of human tissue and the time-intensive nature of acquiring contractility data at multiple loads, we were unable to perform studies using multiple concentrations of ketone or drug. For this reason, we opted to use concentrations at the upper ends of the therapeutic range to determine a saturating effect of drug. Our model is limited in its capacity to study drug effects on end diastolic stress-strain relationship (EDSSR). We found significant differences in EDSSR in control slices following addition of DMSO, confounding our ability to assess impact of other pharmacologic interventions on this parameter. Our model focuses on acute and direct effects of SGLTi and other compounds on human heart tissue and does not capture potential chronic systemic and remodeling effects of SGLTi or ketones. We did not study drug effects that occur over longer time periods (i.e. days-weeks) and were thus unable to assess temporal effects of ketones or SGLTi. It is possible that other systemic and direct effects of SGLTi including metabolic reprogramming or myocardial remodeling have downstream effects on contractility which occur over a longer period.^19,34,60,63^ Current work is aimed at optimizing hLMS culture to enable more biomimetic *ex vivo* conditions. While we found similar effects of SGLTi in NF and HCM tissue, we did not study the effects of SGLTi in failing human hearts. Finally, our work does not identify a clear mechanism or target of SGLT inhibition in the heart. While we see a clear reduction in work following treatment with SGLTi, it is unclear whether this effect is driven by alterations in excitation-contraction coupling, changes in myofilament calcium sensitivity, modifications in myocardial energetics or substrate utilization, or a combination of mechanisms.^54,61^

### Conclusions

Acute contractile phenotyping of human living myocardial slices reflects the *in vivo* state, confirming hypercontractility in HCM and revealing dependence of age, hypertension, ejection fraction and left ventricular mass on contractility. The drug-testing methodology recapitulates known effects of isoproterenol and mavacamten and identifies new disease and load-dependent effects. SGLTi and ketones have distinct, discordant effects on human heart tissue contractility, underscoring the need for future studies to define the mechanisms governing potentially favorable effects of SGLTi in HCM.

## ACKNOWLEDGEMENTS

The authors would like to thank the Penn Metabolomics Core (RRID:SCR_022381) supported by the Penn Cardiovascular Institute and, in part, by NCI P30 CA016520 and NIH P30DK050306.

## References

1. Maron BJ. Clinical Course and Management of Hypertrophic Cardiomyopathy. N Engl J Med. 2018;379:655–668. doi: 10.1056/NEJMra1710575

2. Maron BJ, Maron MS. Hypertrophic cardiomyopathy. Lancet. 2013;381:242–255. doi: 10.1016/S0140-6736(12)60397-3

3. Day SM. Nonobstructive Hypertrophic Cardiomyopathy-The High-Hanging Fruit. JAMA Cardiol. 2019;4:235–236. doi: 10.1001/jamacardio.2018.4953

4. Ommen SR, Mital S, Burke MA, Day SM, Deswal A, Elliott P, Evanovich LL, Hung J, Joglar JA, Kantor P, et al. 2020 AHA/ACC Guideline for the Diagnosis and Treatment of Patients With Hypertrophic Cardiomyopathy: Executive Summary: A Report of the American College of Cardiology/American Heart Association Joint Committee on Clinical Practice Guidelines. Circulation. 2020;142:e533–e557. doi: 10.1161/cir.0000000000000938

5. Olivotto I, Oreziak A, Barriales-Villa R, Abraham TP, Masri A, Garcia-Pavia P, Saberi S, Lakdawala NK, Wheeler MT, Owens A, et al. Mavacamten for treatment of symptomatic obstructive hypertrophic cardiomyopathy (EXPLORER-HCM): a randomised, double-blind, placebo-controlled, phase 3 trial. Lancet. 2020;396:759–769. doi: 10.1016/s0140-6736(20)31792-x

6. Maron MS, Masri A, Nassif ME, Barriales-Villa R, Arad M, Cardim N, Choudhury L, Claggett B, Coats CJ, Dungen HD, et al. Aficamten for Symptomatic Obstructive Hypertrophic Cardiomyopathy. N Engl J Med. 2024;390:1849–1861. doi: 10.1056/NEJMoa2401424

7. Desai MY, Owens AT, Abraham T, Olivotto I, Garcia-Pavia P, Lopes RD, Elliott P, Fernandes F, Verheyen N, Maier L, et al. Mavacamten in Symptomatic Nonobstructive Hypertrophic Cardiomyopathy. New England Journal of Medicine. 2025;393:961–972. doi: doi:10.1056/NEJMoa2505927

8. Maron MS, Rowin EJ, Olivotto I, Casey SA, Arretini A, Tomberli B, Garberich RF, Link MS, Chan RHM, Lesser JR, Maron BJ. Contemporary Natural History and Management of Nonobstructive Hypertrophic Cardiomyopathy. Journal of the American College of Cardiology. 2016;67:1399–1409. doi: 10.1016/j.jacc.2016.01.023

9. Force T, Bonow RO, Houser SR, Solaro RJ, Hershberger RE, Adhikari B, Anderson ME, Boineau R, Byrne BJ, Cappola TP, et al. Research priorities in hypertrophic cardiomyopathy: report of a Working Group of the National Heart, Lung, and Blood Institute. Circulation. 2010;122:1130–1133. doi: 122/11/1130 [pii] 10.1161/CIRCULATIONAHA.110.950089

10. Ghionzoli N, Gentile F, Del Franco AM, Castiglione V, Aimo A, Giannoni A, Burchielli S, Cameli M, Emdin M, Vergaro G. Current and emerging drug targets in heart failure treatment. Heart Fail Rev. 2022;27:1119–1136. doi: 10.1007/s10741-021-10137-2

11. Haghighat L, DeJong C, Teerlink JR. New and future heart failure drugs. Nat Cardiovasc Res. 2024;3:1389–1407. doi: 10.1038/s44161-024-00576-z

12. Fine B, Vunjak-Novakovic G. Shortcomings of Animal Models and the Rise of Engineered Human Cardiac Tissue. ACS Biomater Sci Eng. 2017;3:1884–1897. doi: 10.1021/acsbiomaterials.6b00662

13. Camelliti P, Al-Saud SA, Smolenski RT, Al-Ayoubi S, Bussek A, Wettwer E, Banner NR, Bowles CT, Yacoub MH, Terracciano CM. Adult human heart slices are a multicellular system suitable for electrophysiological and pharmacological studies. J Mol Cell Cardiol. 2011;51:390–398. doi: 10.1016/j.yjmcc.2011.06.018

14. Pitoulis FG, Smith JJ, Pamias-Lopez B, de Tombe PP, Hayman D, Terracciano CM. MyoLoop: Design, development and validation of a standalone bioreactor for pathophysiological electromechanical in vitro cardiac studies. Experimental Physiology.n/a. doi: 10.1113/EP091247

15. Perbellini F, Thum T. Living myocardial slices: a novel multicellular model for cardiac translational research. European Heart Journal. 2019;41:2405–2408. doi: 10.1093/eurheartj/ehz779

16. Fischer C, Milting H, Fein E, Reiser E, Lu K, Seidel T, Schinner C, Schwarzmayr T, Schramm R, Tomasi R, et al. Long-term functional and structural preservation of precision-cut human myocardium under continuous electromechanical stimulation in vitro. Nat Commun. 2019;10:117. doi: 10.1038/s41467-018-08003-1

17. Watson SA, Duff J, Bardi I, Zabielska M, Atanur SS, Jabbour RJ, Simon A, Tomas A, Smolenski RT, Harding SE, et al. Biomimetic electromechanical stimulation to maintain adult myocardial slices in vitro. Nat Commun. 2019;10:2168. doi: 10.1038/s41467-019-10175-3

18. van der Geest JSA, Benavente ED, van Ham WB, Doevendans PA, van Laake LW, de Boer TP, Sampaio-Pinto V, Sluijter JPG. Characterization of human living myocardial slices culture-induced adaptations: a translational perspective. J Mol Cell Cardiol Plus. 2025;13:100465. doi: 10.1016/j.jmccpl.2025.100465

19. Lopaschuk GD, Verma S. Mechanisms of Cardiovascular Benefits of Sodium Glucose Co-Transporter 2 (SGLT2) Inhibitors: A State-of-the-Art Review. JACC Basic Transl Sci. 2020;5:632–644. doi: 10.1016/j.jacbts.2020.02.004

20. Lam CSP, Chandramouli C, Ahooja V, Verma S. SGLT-2 Inhibitors in Heart Failure: Current Management, Unmet Needs, and Therapeutic Prospects. Journal of the American Heart Association. 2019;8:e013389. doi: 10.1161/JAHA.119.013389

21. Verma S, McMurray JJV, Cherney DZI. The Metabolodiuretic Promise of Sodium-Dependent Glucose Cotransporter 2 Inhibition: The Search for the Sweet Spot in Heart Failure. JAMA Cardiology. 2017;2:939–940. doi: 10.1001/jamacardio.2017.1891

22. Cowie MR, Fisher M. SGLT2 inhibitors: mechanisms of cardiovascular benefit beyond glycaemic control. Nature Reviews Cardiology. 2020;17:761–772. doi: 10.1038/s41569-020-0406-8

23. Horton JL, Davidson MT, Kurishima C, Vega RB, Powers JC, Matsuura TR, Petucci C, Lewandowski ED, Crawford PA, Muoio DM, et al. The failing heart utilizes 3-hydroxybutyrate as a metabolic stress defense. JCI Insight. 2024;4. doi: 10.1172/jci.insight.124079

24. Matsuura TR, Puchalska P, Crawford PA, Kelly DP. Ketones and the Heart: Metabolic Principles and Therapeutic Implications. Circ Res. 2023;132:882–898. doi: 10.1161/circresaha.123.321872

25. Nielsen R, Møller N, Gormsen LC, Tolbod LP, Hansson NH, Sorensen J, Harms HJ, Frøkiær J, Eiskjaer H, Jespersen NR, et al. Cardiovascular Effects of Treatment With the Ketone Body 3-Hydroxybutyrate in Chronic Heart Failure Patients. Circulation. 2019;139:2129–2141. doi: 10.1161/circulationaha.118.036459

26. Previs MJ, O’Leary TS, Morley MP, Palmer BM, LeWinter M, Yob JM, Pagani FD, Petucci C, Kim MS, Margulies KB, et al. Defects in the Proteome and Metabolome in Human Hypertrophic Cardiomyopathy. Circ Heart Fail. 2022;15:e009521. doi: 10.1161/circheartfailure.121.009521

27. Selvaraj S, Kelly DP, Margulies KB. Implications of Altered Ketone Metabolism and Therapeutic Ketosis in Heart Failure. Circulation. 2020;141:1800–1812. doi: 10.1161/CIRCULATIONAHA.119.045033

28. McCarthy CG, Chakraborty S, Singh G, Yeoh BS, Schreckenberger ZJ, Singh A, Mell B, Bearss NR, Yang T, Cheng X, et al. Ketone body β-hydroxybutyrate is an autophagy-dependent vasodilator. JCI Insight. 2021;6. doi: 10.1172/jci.insight.149037

29. Lopaschuk GD, Dyck JRB. Ketones and the cardiovascular system. Nature Cardiovascular Research. 2023;2:425-437. doi: 10.1038/s44161-023-00259-1

30. Berg-Hansen K, Gopalasingam N, Christensen KH, Ladefoged B, Andersen MJ, Poulsen SH, Borlaug BA, Nielsen R, Moller N, Wiggers H. Cardiovascular Effects of Oral Ketone Ester Treatment in Patients With Heart Failure With Reduced Ejection Fraction: A Randomized, Controlled, Double-Blind Trial. Circulation. 2024;149:1474–1489. doi: 10.1161/CIRCULATIONAHA.123.067971

31. Gopalasingam N, Berg-Hansen K, Christensen KH, Ladefoged BT, Poulsen SH, Andersen MJ, Borlaug BA, Nielsen R, Møller N, Wiggers H. Randomized Crossover Trial of 2-Week Ketone Ester Treatment in Patients With Type 2 Diabetes and Heart Failure With Preserved Ejection Fraction. Circulation. 2024;150:1570–1583. doi: 10.1161/CIRCULATIONAHA.124.069732

32. Berger JH, Matsuura TR, Bowman CE, Taing R, Patel J, Lai L, Leone TC, Reagan JD, Haldar SM, Arany Z, Kelly DP. SGLT2 Inhibitors Act Independently of SGLT2 to Confer Benefit for HFrEF in Mice. Circulation Research. 2024;135:632–634. doi: 10.1161/CIRCRESAHA.124.324823

33. Philippaert K, Kalyaanamoorthy S, Fatehi M, Long W, Soni S, Byrne NJ, Barr A, Singh J, Wong J, Palechuk T, et al. Cardiac Late Sodium Channel Current Is a Molecular Target for the Sodium/Glucose Cotransporter 2 Inhibitor Empagliflozin. Circulation. 2021;143:2188–2204. doi: 10.1161/CIRCULATIONAHA.121.053350

34. Forelli N, Eaton D, Patel J, Bowman CE, Kawakami R, Kuznetsov IA, Li K, Brady C, Bedi K, Yang Y, et al. SGLT2 inhibitors activate pantothenate kinase in the human heart. bioRxiv. 2024. doi: 10.1101/2024.07.26.605401

35. Pabel S, Wagner S, Bollenberg H, Bengel P, Kovács Á, Schach C, Tirilomis P, Mustroph J, Renner A, Gummert J, et al. Empagliflozin directly improves diastolic function in human heart failure. Eur J Heart Fail. 2018;20:1690–1700. doi: 10.1002/ejhf.1328

36. Amesz JH, Langmuur SJJ, Epskamp N, Bogers A, de Groot NMS, Manintveld OC, Taverne Y. Acute Biomechanical Effects of Empagliflozin on Living Isolated Human Heart Failure Myocardium. Cardiovasc Drugs Ther. 2023. doi: 10.1007/s10557-023-07434-3

37. Wijnker PJM, Dinani R, van der Laan NC, Algul S, Knollmann BC, Verkerk AO, Remme CA, Zuurbier CJ, Kuster DWD, van der Velden J. Hypertrophic cardiomyopathy dysfunction mimicked in human engineered heart tissue and improved by sodium-glucose cotransporter 2 inhibitors. Cardiovasc Res. 2024;120:301–317. doi: 10.1093/cvr/cvae004

38. van der Geest JSA, de Boer TP, Terracciano CM, Thum T, Dendorfer A, Doevendans PA, van Laake LW, Sluijter JPG, Sampaio-Pinto V. Living myocardial slices: walking the path towards standardization. Cardiovasc Res. 2025;121:1011–1023. doi: 10.1093/cvr/cvaf079

39. Pitoulis FG, Watson SA, Perbellini F, Terracciano CM. Myocardial slices come to age: an intermediate complexity in vitro cardiac model for translational research. Cardiovasc Res. 2020;116:1275–1287. doi: 10.1093/cvr/cvz341

40. Pitoulis FG, Nunez-Toldra R, Xiao K, Kit-Anan W, Mitzka S, Jabbour RJ, Harding SE, Perbellini F, Thum T, de Tombe PP, Terracciano CM. Remodelling of adult cardiac tissue subjected to physiological and pathological mechanical load in vitro. Cardiovasc Res. 2022;118:814–827. doi: 10.1093/cvr/cvab084

41. Dolder FWvd, Dinani R, Warnaar VAJ, Vučković S, Passadouro AS, Nassar AA, Ramsaroep AX, Burchell GB, Schoonmade LJ, Velden Jvd, Goversen B. Experimental Models of Hypertrophic Cardiomyopathy. JACC: Basic to Translational Science. 2025;10:511–546. doi: doi:10.1016/j.jacbts.2024.10.017

42. Nassar A, Warnaar VAJ, Duursma I, Schomakers BV, Chaami C, Ochala J, van Weeghel M, Houtkooper RH, Michels M, Dendorfer A, et al. Ex-vivo culture of human hypertrophic cardiomyopathy hearts: functional and metabolic changes during long-term culture. iScience. doi: 10.1016/j.isci.2026.115308

43. Sequeira V, Maack C, Reil G-H, Reil J-C. Exploring the Connection Between Relaxed Myosin States and the Anrep Effect. Circulation Research. 2024;134:117–134. doi: 10.1161/CIRCRESAHA.123.323173

44. Reil J-C, Sequeira V, Reil G-H, Scholtz S, Rudolph V, Maack C, Serruys P, Steendijk P. Investigating the Anrep Effect in Hypertrophic Obstructive Cardiomyopathy With Invasive Pressure-Volume Analysis. JACC: Advances. 2025;4:101728. doi: 10.1016/j.jacadv.2025.101728

45. Fleg JL, Strait J. Age-associated changes in cardiovascular structure and function: a fertile milieu for future disease. Heart Fail Rev. 2012;17:545–554. doi: 10.1007/s10741-011-9270-2

46. Selvaraj S, Hu R, Vidula MK, Dugyala S, Tierney A, Ky B, Margulies KB, Shah SH, Kelly DP, Bravo PE. Acute Echocardiographic Effects of Exogenous Ketone Administration in Healthy Participants. J Am Soc Echocardiogr. 2022;35:305–311. doi: 10.1016/j.echo.2021.10.017

47. Vite A, Matsuura TR, Bedi KC, Flam EL, Arany Z, Kelly DP, Margulies KB. Functional Impact of Alternative Metabolic Substrates in Failing Human Cardiomyocytes. JACC Basic Transl Sci. 2024;9:1–15. doi: 10.1016/j.jacbts.2023.07.009

48. Edgar EN, Vasco S, Christoph M, Julien O, Jolanda vdV. Metabolic alterations in human hypertrophic cardiomyopathy. The Journal of Cardiovascular Aging. 2025;5:11.

49. Toepfer CN, Garfinkel AC, Venturini G, Wakimoto H, Repetti G, Alamo L, Sharma A, Agarwal R, Ewoldt JK, Cloonan P, et al. Myosin Sequestration Regulates Sarcomere Function, Cardiomyocyte Energetics, and Metabolism, Informing the Pathogenesis of Hypertrophic Cardiomyopathy. Circulation. 2020;141:828–842. doi: 10.1161/CIRCULATIONAHA.119.042339

50. Day SM, Tardiff JC, Ostap EM. Myosin modulators: emerging approaches for the treatment of cardiomyopathies and heart failure. The Journal of clinical investigation. 2022;132. doi: 10.1172/JCI148557

51. Ochala J, Feng M, Wang Q, Chaami C, Nollet E, Lewis CTA, Hessel AL, Michels M, Bedi KC, Jr., Margulies KB, et al. Heterogeneous Dysregulation of Myosin Super-Relaxation and Energetics in Hypertrophic Cardiomyopathy. Circ Heart Fail. 2025:e012614. doi: 10.1161/CIRCHEARTFAILURE.124.012614

52. Nyitrai M, Geeves MA. Adenosine diphosphate and strain sensitivity in myosin motors. Philos Trans R Soc Lond B Biol Sci. 2004;359:1867–1877. doi: 10.1098/rstb.2004.1560

53. Irving M. Functional control of myosin motors in the cardiac cycle. Nat Rev Cardiol. 2025;22:9–19. doi: 10.1038/s41569-024-01063-5

54. Pabel S, Wagner S, Bollenberg H, Bengel P, Kovacs A, Schach C, Tirilomis P, Mustroph J, Renner A, Gummert J, et al. Empagliflozin directly improves diastolic function in human heart failure. Eur J Heart Fail. 2018;20:1690–1700. doi: 10.1002/ejhf.1328

55. Coppini R, Ferrantini C, Yao L, Fan P, Del Lungo M, Stillitano F, Sartiani L, Tosi B, Suffredini S, Tesi C, et al. Late sodium current inhibition reverses electromechanical dysfunction in human hypertrophic cardiomyopathy. Circulation. 2013;127:575–584. doi: 10.1161/circulationaha.112.134932

56. A Randomized, Double-blind, Placebo-controlled, Parallel-group, Multicenter Study to Evaluate the Efficacy and Safety of SOtaglifloziN in symptomATic Obstructive And Non-obstructive Hypertrophic CardioMyopathy (SONATA-HCM). In; 2024.

57. Sherratt SC, Weindorf AV, Libby P, Bhatt DL, Mason P. Abstract 16947: Sotagliflozin, a Dual SGLT 1 and 2 Inhibitor, Has High Membrane Affinity and Hydrocarbon Core Location Compared With Empagliflozin. Circulation. 2023;148:A16947–A16947. doi: doi:10.1161/circ.148.suppl_1.16947

58. Banerjee SK, McGaffin KR, Pastor-Soler NM, Ahmad F. SGLT1 is a novel cardiac glucose transporter that is perturbed in disease states. Cardiovasc Res. 2009;84:111–118. doi: 10.1093/cvr/cvp190

59. Saucedo-Orozco H, Voorrips SN, Yurista SR, de Boer RA, Westenbrink BD. SGLT2 Inhibitors and Ketone Metabolism in Heart Failure. J Lipid Atheroscler. 2022;11:1–19. doi: 10.12997/jla.2022.11.1.1

60. Goedeke L, Ma Y, Gaspar RC, Nasiri A, Lee J, Zhang D, Galsgaard KD, Hu X, Zhang J, Guerrera N, et al. SGLT2 inhibition alters substrate utilization and mitochondrial redox in healthy and failing rat hearts. J Clin Invest. 2024;134. doi: 10.1172/JCI176708

61. Berger JH, Finck BN. Beyond ketosis: the search for the mechanism underlying SGLT2-inhibitor benefit continues. J Clin Invest. 2024;134. doi: 10.1172/JCI187097

62. Vissing CR, Axelsson Raja A, Helms AS, Saberi S, Owens AT, Rossano JW, Abrams DJ, Ingles J, Gray B, Lampert R, et al. Differences in Disease Trajectory, Comorbidities, and Mortality in Sarcomeric and Nonsarcomeric Hypertrophic Cardiomyopathy. Circulation. 2026. doi: 10.1161/CIRCULATIONAHA.125.076036

63. Santos-Gallego CG, Requena-Ibanez JA, San Antonio R, Ishikawa K, Watanabe S, Picatoste B, Flores E, Garcia-Ropero A, Sanz J, Hajjar RJ, et al. Empagliflozin Ameliorates Adverse Left Ventricular Remodeling in Nondiabetic Heart Failure by Enhancing Myocardial Energetics. J Am Coll Cardiol. 2019;73:1931–1944. doi: 10.1016/j.jacc.2019.01.056

