## Supplementary Figures and Tables for "Load-Dependent Effects of Sodium Glucose Co-Transporter Inhibitors on Work in Human Hypertrophic Cardiomyopathy Living Myocardial Slices"

### SUPPLEMENTAL METHODS

**Human Myocardial Tissue Procurement:** Human myocardial tissue was procured under the protocols and ethical regulations approved by the Institutional Review Boards at the University of Pennsylvania and the Gift-of-Life Donor Program (Philadelphia, PA). Non-failing hearts were obtained at the time of organ donation and were arrested *in situ* using ice-cold cardioplegia as previously described.[1] A 1 cm x 1 cm full-thickness portion of either the left ventricular free wall or septum was dissected and saved for downstream use. HCM myocardial tissue, obtained at the time of septal reduction surgery, was placed immediately into ice-cold cardioplegia, and transferred to the lab.

**Generation of Left Ventricular Living Myocardial Slices:** Within 30 minutes of explant, the tissue was transferred to oxygen-bubbled ice-cold Slicing Solution (140 mM NaCl, 5.4 mM KCl, 10 mM HEPES, 10 mM D-glucose, 1 mM $CaCl_2$, 0.6 mM $MgCl_2$, 3 g/L 2,3-butanedione monoxime (BDM), pH 7.4). Excess tissue was trimmed and tissue block adhered to the metal puck of the Campden Instruments 7000 smz2 Vibratome using N-butyl cyanoacrylate tissue adhesive (3M). Tissue blocks were sliced using the following parameters: 2 mm amplitude, 80 Hz, 0.03 mm/s advance speed. Slices were trimmed to 5 mm x 5 mm or 8 mm x 8 mm and adhered using tissue adhesive to custom laser cut (Epilog Fusion Pro 36) anchors (0.2 mm thick Delrin/Acetal (McMaster-Carr)) (**Figure S1**). Up to 7 300 µm slices were analyzed per heart (**Table S1a-b**). Slices were then placed into a Recording Solution (140 mM NaCl, 2.7 mM KCl, 10 mM HEPES, 10 mM D-glucose, 1.8 mM $CaCl_2$, 0.6 mM $MgCl_2$, pH 7.4) with or without supplemental ketones, 6 mM 3-OHB ((R)-3-Hydroxybutyric acid, Sigma) and allowed to equilibrate for one hour in an incubator (37°C, 5% $CO_2$).

**Contractility Measurement and Drug Testing of hLMS:** Following equilibration, anchor stays were cut, and slices were mounted to the IonOptix Cardiac Slice System in recirculating

Recording Solution at 37°C with bubbled 100% oxygen. Following an additional ten-minute equilibration period at slack length (0% strain), baseline mechanics were measured. Tissues were stretched to 4% strain and paced at 1 Hz to generate isometric force transients. Biomimetic work loops were generated consisting of four phases reflecting valve opening and closure in intact hearts: (1) an isometric contraction phase starting immediately following paced excitation where hLMS were held at fixed length, (2) a shortening phase where at a predesignated developed force (*afterload*), hLMS are allowed to shorten proportional to their contractile force, (3) an isometric relaxation phase where hLMS are held in a shortened position as force falls, and (4) a diastolic lengthening phase where hLMS are returned to a predesignated resting length prior to the initiation of the next cycle. Work loops were generated at afterloads of 25%, 50%, and 75% of developed isometric force. Measurements were repeated at 8%, 12%, and 16% strain.

Following baseline contractility measurements, hLMS were de-stretched to 0% strain. Drug or an equivalent volume of dimethyl sulfoxide (DMSO, Corning) as a vehicle control was added to Recording Solution and allowed to recirculate for 20 minutes. After drug treatment, the same mechanical protocol was performed. Drugs used in this study include: 5 μM isoproterenol (Sigma), 1 μM mavacamten (Medchem Express), 10 μM sotagliflozin (Lexicon Pharmaceuticals) or 10 μM empagliflozin (Cayman Chemical Company).

**Work Loop Data Analysis:** Data were processed and analyzed using IonWizard (IonOptix). Raw force data were normalized to stress (force per cross-sectional area of a given slice) and length data normalized to strain (% stretch from slack length). Stress-strain work loops were visualized and analyzed. For each set of work loops, 5 representative loops from each load were processed and data parameters averaged. Analyzed parameters included stress ($mN/mm^2$) and strain (%) at each corner of the work loop (i.e. at the transition from isometric to shortening/lengthening phases or vice versa), duration of time spent in each of the 4 phases of the work loop (ms), the maximum and minimum stress of the work loop ($mN/mm^2$), developed stress (systolic rise in stress, $mN/mm^2$), stroke strain (end-diastolic – end-systolic strain (%)), and work (area within the work loop, $\mu J/mm^2$). Slopes of the end systolic stress strain and end diastolic stress strain relationships ($mN/mm^2\%$) were also calculated.

To perform principal components analysis (PCA) on baseline data, all measured and calculated parameters in the baseline set of work loops were used and classified according to underlying disease (NF or HCM) or fuel composition (glucose only or glucose supplemented with ketones). Each PCA point reflects a single slice. For drug effect PCAs, the fold change of all parameters was determined between the post-drug set of work loops and baseline set, then PCA calculated from the fold change. Each dataset was analyzed with the FactoMineR package (version 2.12). PCA was carried out on centered and scaled variables using the PCA function with default settings. The eigenvalues were extracted to quantify the percentage of variance explained by each principal component. Individual coordinates from the PCA were exported and visualized with ggplot2 (version 3.5.2). Scatterplots were constructed using the first two principal components, and samples were color-coded according to the grouping variable of interest. Ninety-five percent confidence ellipses were generated around each group using the stat_ellipse function to assess clustering and separation among groups. Each dot represents one slice.

**Linear Regression Modeling and Statistics:** Statistical analysis was conducted in STATA (StataNow/SE 18.5). Mixed effect linear regression modeling was used to examine associations between clinical parameters (disease state (NF vs. HCM), age, sex, hypertension, diabetes, body mass index), experimental conditions (fuel type, drug type), and contractility parameters (work, developed stress, or end-diastolic stress-strain slope). Models were fitted separately for NF and HCM slices, and at each of 3 afterloads. All models included fixed effects for age (continuous, centered), sex, body mass index (BMI, centered), diabetes (yes or no), hypertension (yes or no), left ventricular ejection fraction (%, continuous, centered), and left ventricular mass index (LVMI, $g/m^2$, continuous, centered), interaction terms between each clinical parameter and strain (centered), and random intercepts for heart and slice to account for non-independence of repeated measures within hearts and within myocardial slices. Interaction terms between strain and variables of interest were included to assess association between a given variable and tissue responsiveness to increasing strain.

Parameter estimates are reported as regression coefficients with corresponding standard errors and two-sided p-values. Average marginal effects and pairwise contrasts were obtained using the *margins* and *pwcompare(effect)* commands in STATA. Where relevant, *lincom* was used to generate custom contrasts of regression slopes across experimental conditions. Model outputs are summarized by two parameters: an average at centered strain (i.e. the modeled work or developed stress at 10% strain), and the interaction between the contractile parameter of interest and strain (i.e. change in work per unit increase in strain; work-strain slope or the end diastolic stress strain slope).

**Metabolomics:** Following drug test and mechanics assessment, slices were immediately cut from the anchors, flash-frozen in liquid nitrogen, and stored in -80°C for further processing. Targeted metabolomics were performed by the University of Pennsylvania Metabolomics Core. Briefly, slices were lyophilized overnight and approximately 2.5 mg was homogenized in 250 μL of 50% acidified acetonitrile (0.3% formic acid) for targeted LC/MS/MS metabolomics (acylcarnitines, amino acids, organic acids) according to validated, optimized protocols in our previously published study.[2] Separate aliquots of homogenates were extracted with solvents for each class of metabolites, and then each class was analyzed with a unique LC/MS/MS method to optimize their chromatographic resolution and sensitivity. Quantitation of metabolites in each assay module was achieved using multiple reaction monitoring of calibration solutions and study samples with isotopically labelled internal standards on an Agilent 1290 Infinity UHPLC/6495 triple quadrupole mass spectrometer. Raw data were processed using Mass Hunter quantitative analysis software (Agilent). Calibration curves ($R^2$ = 0.99 or greater) were fitted with either a linear or a quadratic curve with a 1/X or $1/X^2$ weighting.

**SUPPLEMENTAL FIGURES:**

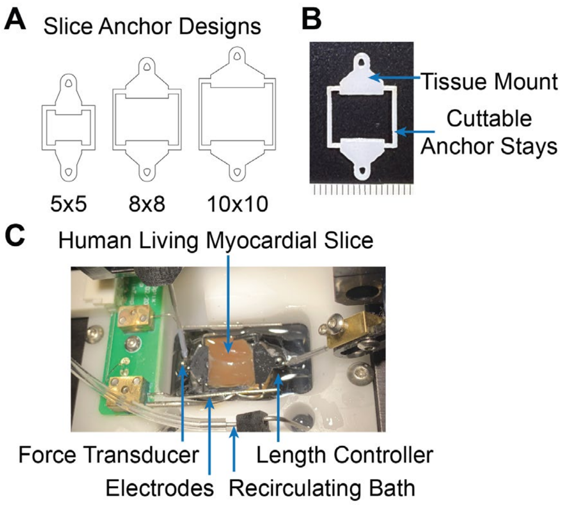

**Figure S1. Slice Anchor Designs. A.** Slice anchor designs were generated at 5-, 8-, and 10-mm lengths and widths to accommodate slices of different sizes. When able, 8 mm x 8 mm slices were used. **B.** Photograph of laser-cut LMS anchor consisting of a tissue mount and anchor stays that were cut prior to mounting. Hashes=1 mm. **C**. Mounted human LMS on anchors fit on mechanical testing rig between a force transducer and length controller capable of generating work loops.

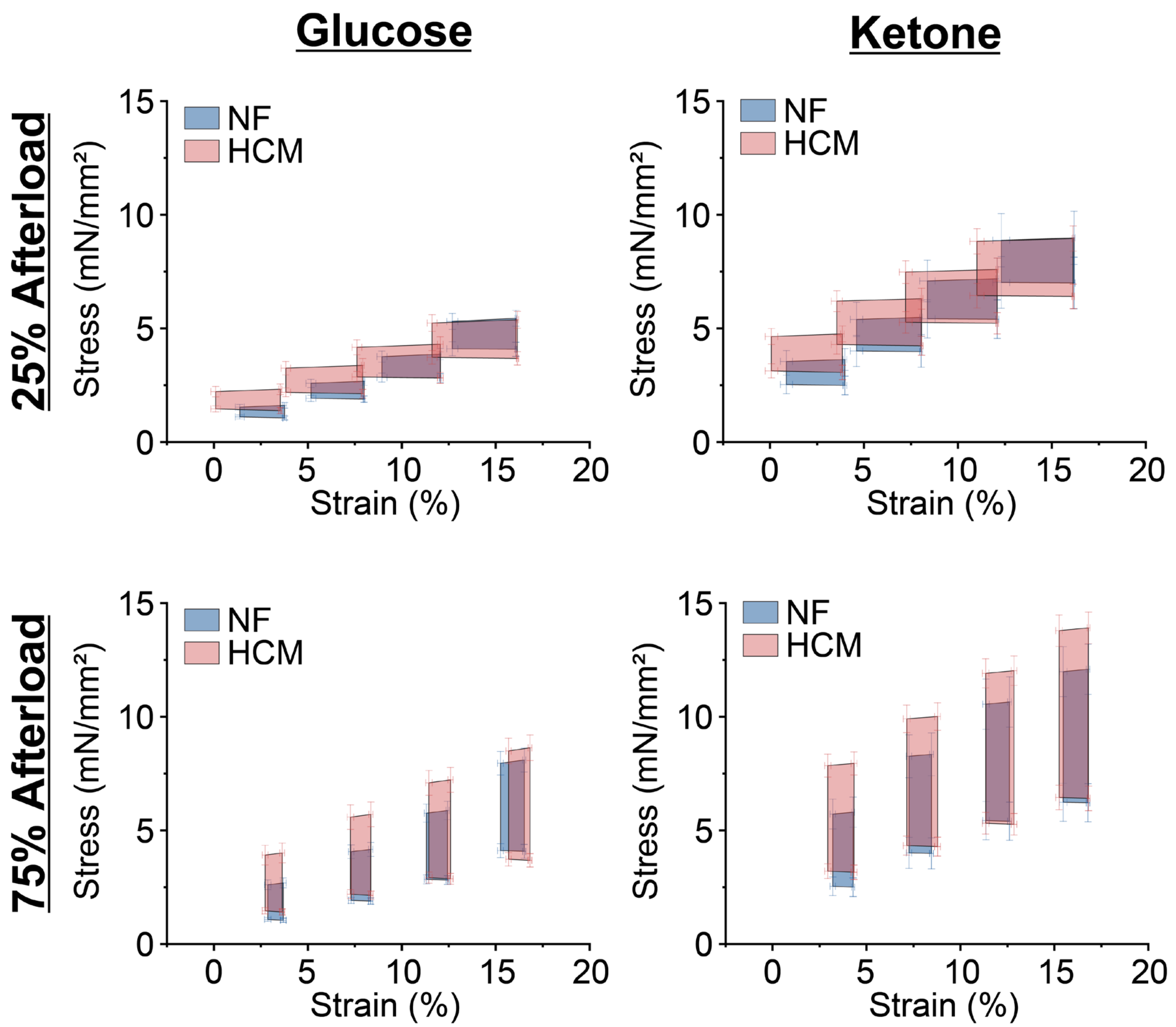

**Figure S2. Average Baseline Work Loops**. Average work loops ± SEM at all preloads and 25% or 75% afterload. Replicates: N=32 hearts, n=120 slices.

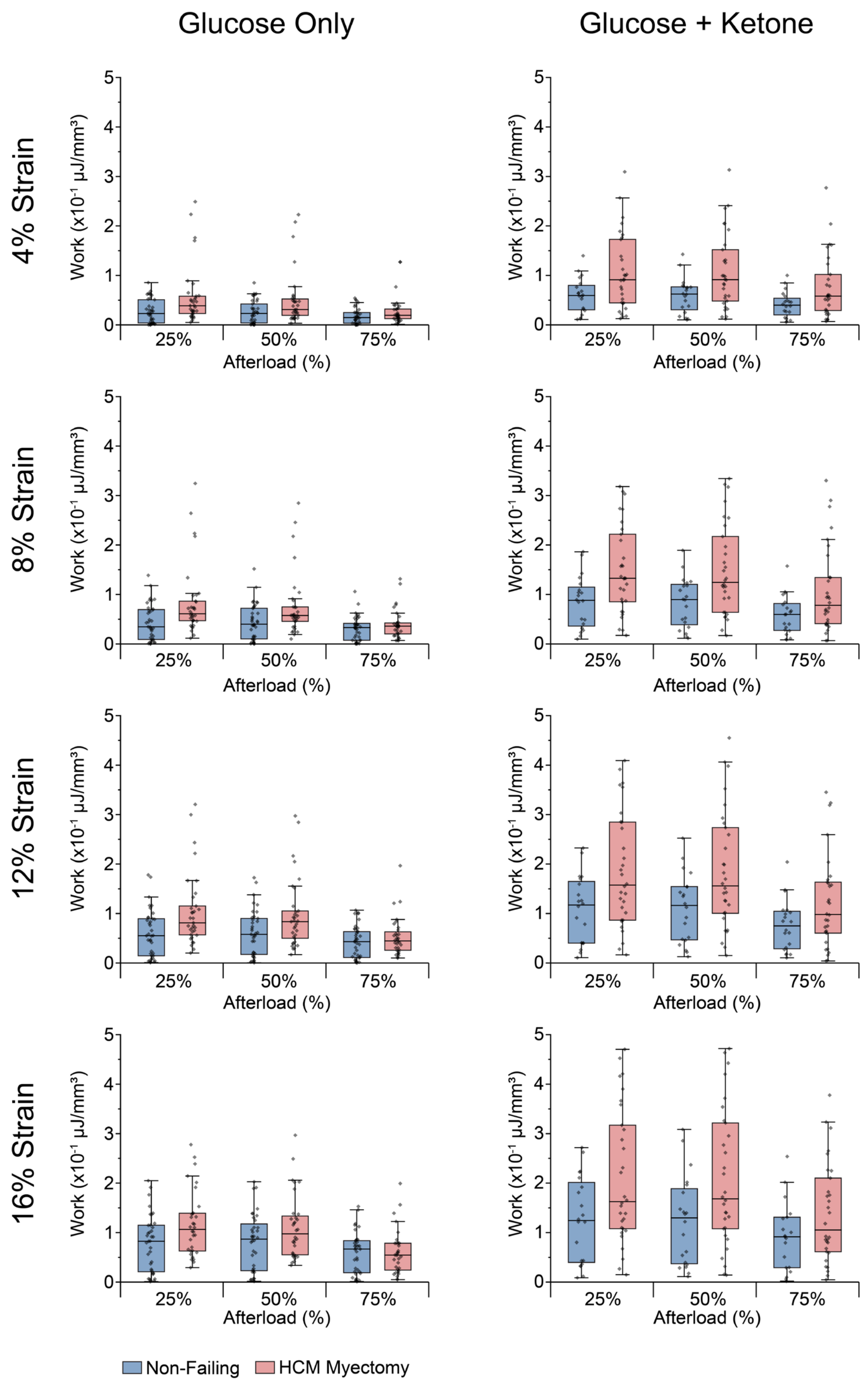

**Figure S3. Baseline Work**. Calculated work at all strains for each slice at 25, 50, and 75% afterloads. Boxes denote median, 25th and 75th percentile and whiskers represent 1.5 interquartile range (IQR). Replicates: N=32 hearts, n=120 slices.

| | 25% Afterload | | 50% Afterload | | 75% Afterload | |
|---|---|---|---|---|---|---|
| Condition | Dev. Stress (SEM) mN/mm$^2$ | p-value vs NF | Dev. Stress (SEM) mN/mm$^2$ | p-value vs NF | Dev. Stress (SEM) mN/mm$^2$ | p-value vs NF |
| Glucose | | | | | | |
| NF | 1.62 (0.31) | | 2.35 (0.46) | | 3.09 (0.59) | |
| HCM | 2.75 (0.31) | **0.027** | 3.99 (0.46) | **0.030** | 5.12 (0.60) | **0.037** |
| Ketone | | | | | | |
| NF | 2.29 (0.41) | | 3.35 (0.60) | | 4.42 (0.79) | |
| HCM | 3.15 (0.34) | 0.17 | 4.68 (0.51) | 0.15 | 6.07 (0.67) | 0.17 |

**Table S1. Baseline Mean Centered Developed Active Stress (at 10% Strain)**. P-values show comparison between HCM and NF in glucose or ketone. P values less than 0.05 are bolded. Replicates: N=32 hearts, n=120 slices. Statistics: Mixed-effects linear regression models with random intercepts for heart and slice.

| | Non-Failing | | HCM | |
|---|---|---|---|---|
| Metabolite | Glucose pmol/mg (CI) | Ketone pmol/mg (CI) | Glucose pmol/mg (CI) | Ketone pmol/mg (CI) |
| 3-HBA | 273.8 (178, 369.59) | 23193.74 (18134.77, 28252.71) | 250.88 (121.3, 380.46) | 24474.56 (18535.71, 30413.41) |
| alpha-KG Acid | 54.07 (35.26, 72.88) | 82.44 (-51.49, 216.37) | 42.45 (32.82, 52.08) | 73.46 (40.96, 105.97) |
| Citrate | 668.66 (457.27, 880.05) | 904.18 (-397.47, 2205.83) | 1094.68 (448.46, 1740.9) | 843.81 (509.21, 1178.4) |
| Fumarate | 23.58 (7.42, 39.73) | 54.39 (14.69, 94.09) | 15.3 (1.63, 28.96) | 33.09 (18.02, 48.16) |
| Lactate | 5537.35 (2267.94, 8806.76) | 6169.87 (1847.1, 10492.65) | 4726.49 (-550.76, 10003.74) | 3318.53 (1899.58, 4737.48) |
| Malate | 222.49 (157.4, 287.59) | 261.44 (154.58, 368.29) | 196.89 (111.6, 282.18) | 160.25 (101.93, 218.56) |
| Pyruvate | 163.34 (91.19, 235.48) | 163.31 (44.42, 282.21) | 107.45 (63.28, 151.61) | 161.1 (99.89, 222.3) |
| Succinate | 71.45 (10.42, 132.48) | 51.65 (12.97, 90.34) | 75.62 (-52.23, 203.46) | 28.46 (16.49, 40.42) |
| C02 | 1669.52 (1289.52, 2049.51) | 1226.63 (638.83, 1814.42) | 1238.37 (502.59, 1974.15) | 1521.61 (876.61, 2166.61) |
| C03 | 9.64 (4.75, 14.52) | 3.49 (1.27, 5.71) | 4.32 (1.64, 6.99) | 2.39 (1.35, 3.43) |
| C03-DC | 0.67 (0.18, 1.17) | 0.91 (0.38, 1.45) | 0.41 (0.29, 0.53) | 0.81 (0.53, 1.09) |
| C04 Butyryl | 7.68 (3.44, 11.92) | 4.84 (-4.09, 13.76) | 8.12 (-2.81, 19.06) | 2.86 (1.28, 4.43) |
| C04 Isobutyryl | 2.14 (0.67, 3.61) | 0.54 (0.04, 1.04) | 9.79 (-5.39, 24.97) | 0.65 (-0.01, 1.3) |
| C04-DC MeMal | 0.46 (0.33, 0.59) | 0.54 (0.07, 1) | 0.4 (0.28, 0.52) | 0.42 (0.24, 0.6) |
| C04-DC Succinyl | 1.52 (0.37, 2.67) | 3.1 (-2.67, 8.87) | 0.62 (0.24, 1.01) | 0.49 (0.25, 0.72) |
| C04-OH Butyryl | 11.57 (6.28, 16.86) | 86.99 (50.54, 123.45) | 7.49 (2.32, 12.67) | 43.97 (9.27, 78.66) |
| C04-OH Isobutyryl | 56.73 (17.85, 95.6) | 65.62 (-25.59, 156.83) | 24.18 (-2.4, 50.75) | 21.47 (9.27, 33.67) |
| C05 2-Methylbutyryl | 2.74 (0.67, 4.8) | 2.21 (0.37, 4.04) | 1.26 (0.64, 1.89) | 1.16 (0.52, 1.8) |
| C05 Isovaleryl | 1.94 (1.28, 2.6) | 0.67 (0.53, 0.82) | 2.55 (-0.48, 5.58) | 1.3 (0.25, 2.35) |
| C05:1 | 3.87 (1.21, 6.53) | 1.66 (0.54, 2.77) | 1.92 (1.31, 2.53) | 1.23 (0.23, 2.22) |
| C05-DC | 0.62 (0.31, 0.93) | 0.65 (0.01, 1.29) | 0.43 (0.2, 0.67) | 0.42 (0.15, 0.68) |
| C05-OH | 15.56 (3.87, 27.25) | 6.51 (1.29, 11.72) | 8.5 (4.93, 12.07) | 4.08 (2.41, 5.74) |
| C06 | 1.17 (0.49, 1.85) | 0.67 (0.04, 1.29) | 0.61 (0.1, 1.12) | 0.31 (0.19, 0.43) |
| C06-OH | 1.24 (0.94, 1.55) | 1.82 (0.2, 3.43) | 1.66 (0.88, 2.44) | 2.25 (0.4, 4.11) |
| C08 | 0.42 (0.07, 0.77) | 0.25 (0.02, 0.47) | 0.25 (0.09, 0.41) | 0.16 (0.11, 0.22) |
| C08:1-OH | 0.18 (0.03, 0.33) | 0.85 (0.02, 1.68) | 0.57 (0.36, 0.78) | 2.38 (-0.07, 4.83) |
| C08-OH | 0.71 (0.26, 1.16) | 0.7 (-0.18, 1.59) | 0.23 (-0.02, 0.48) | 0.27 (0.1, 0.43) |
| C10 | 1.86 (-0.25, 3.97) | 4.29 (2.85, 5.73) | 0.8 (-0.62, 2.22) | 4.06 (3.44, 4.69) |
| C10-OH | 0.47 (-0.09, 1.02) | 0.55 (-0.38, 1.47) | 0.07 (0, 0.14) | 0.15 (0.06, 0.25) |
| C12 | 2.81 (0.76, 4.86) | 1.11 (-0.27, 2.48) | 2.69 (0.84, 4.54) | 0.73 (0.5, 0.96) |
| C12:1 | 0.86 (0.2, 1.53) | 0.42 (-0.2, 1.05) | 0.5 (0.27, 0.73) | 0.21 (0.14, 0.27) |
| C12-OH | 1.11 (0.29, 1.93) | 0.8 (-0.56, 2.17) | 0.51 (0.22, 0.79) | 0.25 (0.12, 0.37) |
| C14 | 4.49 (-0.16, 9.14) | 1.37 (-0.62, 3.36) | 6.46 (1.42, 11.49) | 0.97 (0.52, 1.42) |
| C14:1 | 3.39 (-0.23, 7.01) | 1.19 (-0.43, 2.82) | 2.23 (0.5, 3.95) | 0.56 (0.27, 0.85) |

| | | | | |
|---|---|---|---|---|
| C14:1-OH | 0.4 (0.09, 0.72) | 0.33 (-0.37, 1.04) | 0.17 (0.06, 0.29) | 0.06 (-0.01, 0.13) |
| C14:2 | 1.1 (0.43, 1.76) | 0.64 (-0.57, 1.86) | 0.83 (0.3, 1.36) | 0.22 (0.11, 0.33) |
| C14:2-OH | 0.12 (0.08, 0.15) | 0.11 (-0.24, 0.45) | 0.13 (0.07, 0.2) | 0.03 (-0.28, 0.35) |
| C14-OH | 1.03 (0.26, 1.8) | 0.95 (-0.95, 2.86) | 0.8 (0.32, 1.29) | 0.17 (0.05, 0.29) |
| C16 | 19.26 (0.99, 37.53) | 6.32 (-0.59, 13.22) | 32.23 (7.56, 56.9) | 6.3 (3.98, 8.62) |
| C16:1 | 7.03 (-1.91, 15.97) | 1.54 (-0.89, 3.97) | 10.61 (0.83, 20.4) | 1.02 (0.49, 1.55) |
| C16:1-OH | 1.33 (0.01, 2.65) | 0.96 (-0.89, 2.82) | 1.28 (0.38, 2.19) | 0.22 (0.09, 0.35) |
| C16:2 | 0.91 (0.09, 1.73) | 0.33 (-0.11, 0.78) | 3.2 (0.35, 6.05) | 0.2 (0.08, 0.31) |
| C16:2-OH | 0.37 (0.16, 0.58) | 0.29 (-0.07, 0.65) | 0.33 (0.13, 0.53) | 0.13 (0.07, 0.19) |
| C16-OH | 2.01 (0.2, 3.81) | 1.2 (-0.94, 3.34) | 3.64 (0.52, 6.77) | 0.44 (0.16, 0.72) |
| C18 | 14.64 (-0.86, 30.14) | 5.24 (-1.15, 11.63) | 18.12 (3.92, 32.33) | 5.23 (3.45, 7.01) |
| C18:1 | 83.64 (-51.13, 218.41) | 25 (-9.72, 59.72) | 73.65 (6.19, 141.12) | 13.79 (6.8, 20.78) |
| C18:1-OH | 7.01 (-2.41, 16.42) | 3.45 (-2.96, 9.87) | 6.86 (0.9, 12.81) | 0.73 (-0.07, 1.54) |
| C18:2 | 25.38 (-9.52, 60.28) | 10.3 (-2.53, 23.14) | 34.47 (5.21, 63.73) | 6.2 (2.37, 10.04) |
| C18:2-OH | 1.5 (-0.3, 3.3) | 1.12 (-1.36, 3.59) | 3.01 (0.59, 5.43) | 0.38 (-0.28, 1.04) |
| C18-OH | 1.88 (-0.6, 4.36) | 0.95 (-1.05, 2.95) | 2.35 (0.3, 4.4) | 0.36 (-0.06, 0.78) |
| C20 | 0.78 (-0.27, 1.83) | 0.04 (-0.4, 0.48) | 1.99 (-0.21, 4.19) | 0.74 (-7.43, 8.91) |

**Table S2. Targeted Metabolomics of LMS in Glucose vs. Ketone supplemented Solutions.** Mean amount of select metabolites (pmol) relative to tissue dry mass with 95% Confidence Intervals (CI). Replicates: N=32 hearts; n=8 (NF, glucose), 4 (NF, ketone), 6 (HCM, glucose), 7 (HCM, ketone) slices.

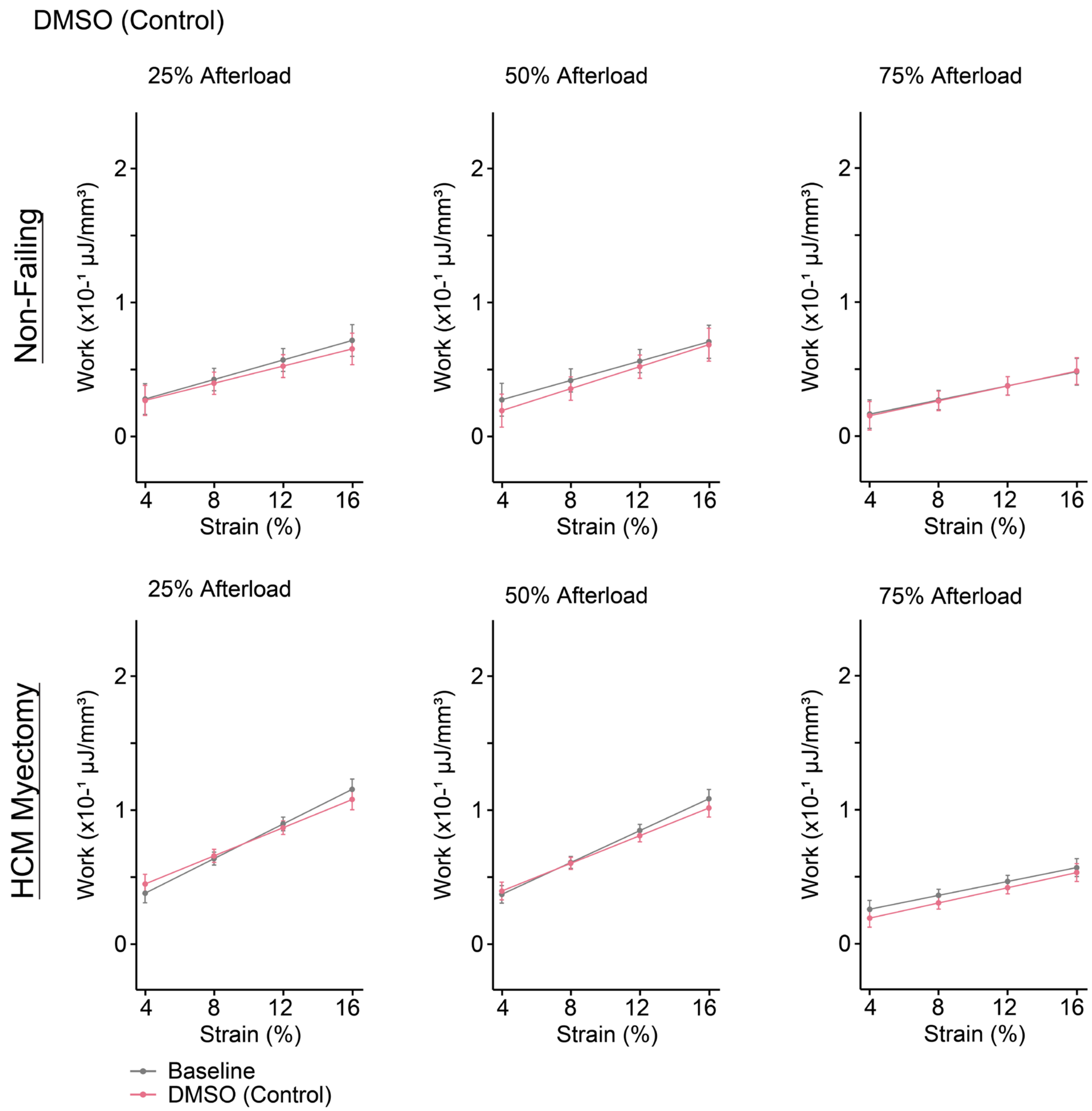

**Figure S4. Mixed Effect Linear Model Output in Response to DMSO.** Mean ± standard error at baseline (gray) and following DMSO administration (pink). Replicates: N=15 hearts; n=15 (DMSO) slices. Statistics: Mixed-effects linear regression models with random intercepts for heart and slice.

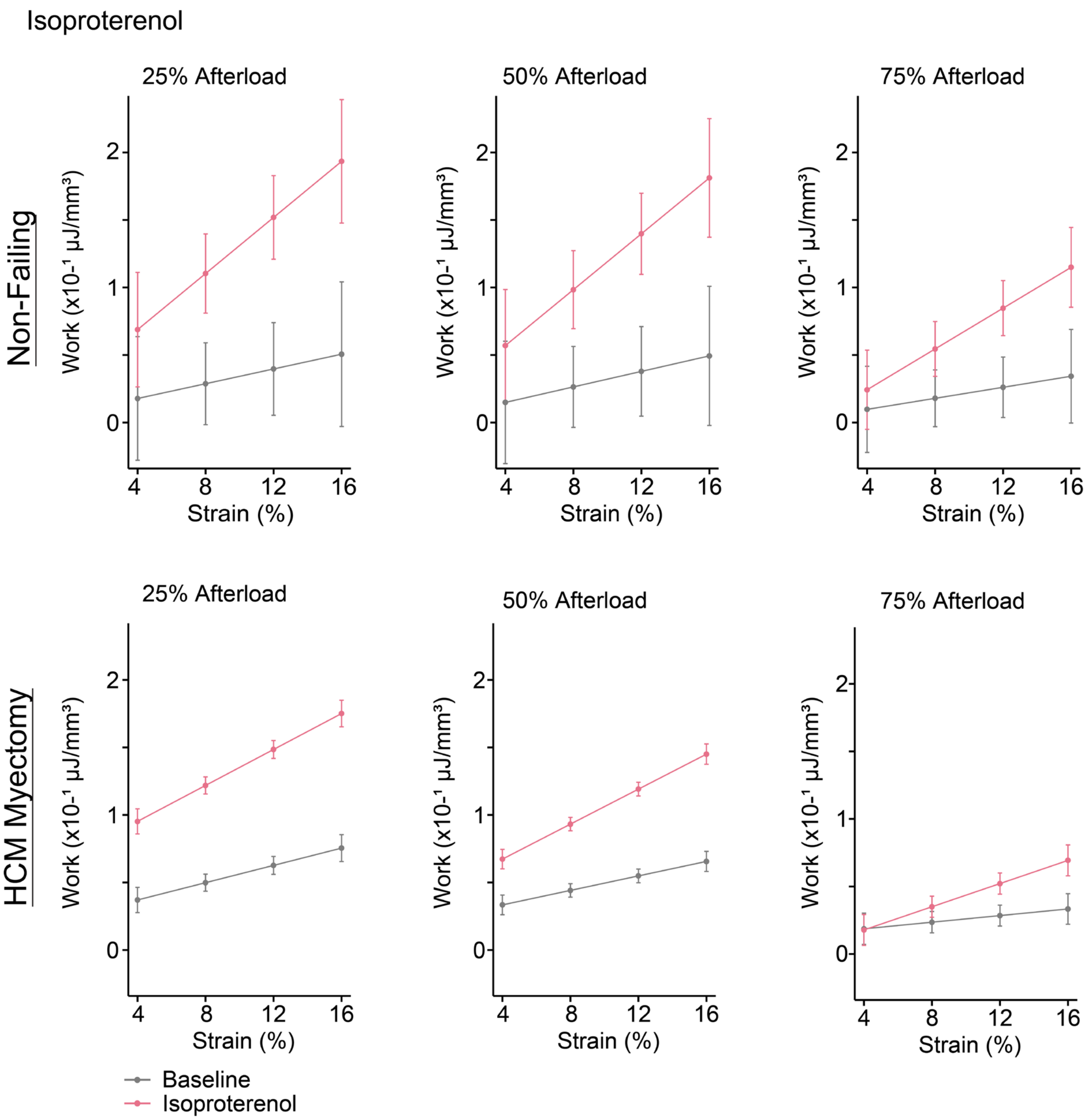

**Figure S5. Mixed Effect Linear Model Output in Response to Isoproterenol.** Mean ± standard error at baseline (gray) and following isoproterenol administration (pink). Replicates: N=6 hearts; 6 (isoproterenol) slices. Statistics: Mixed-effects linear regression models with random intercepts for heart and slice.

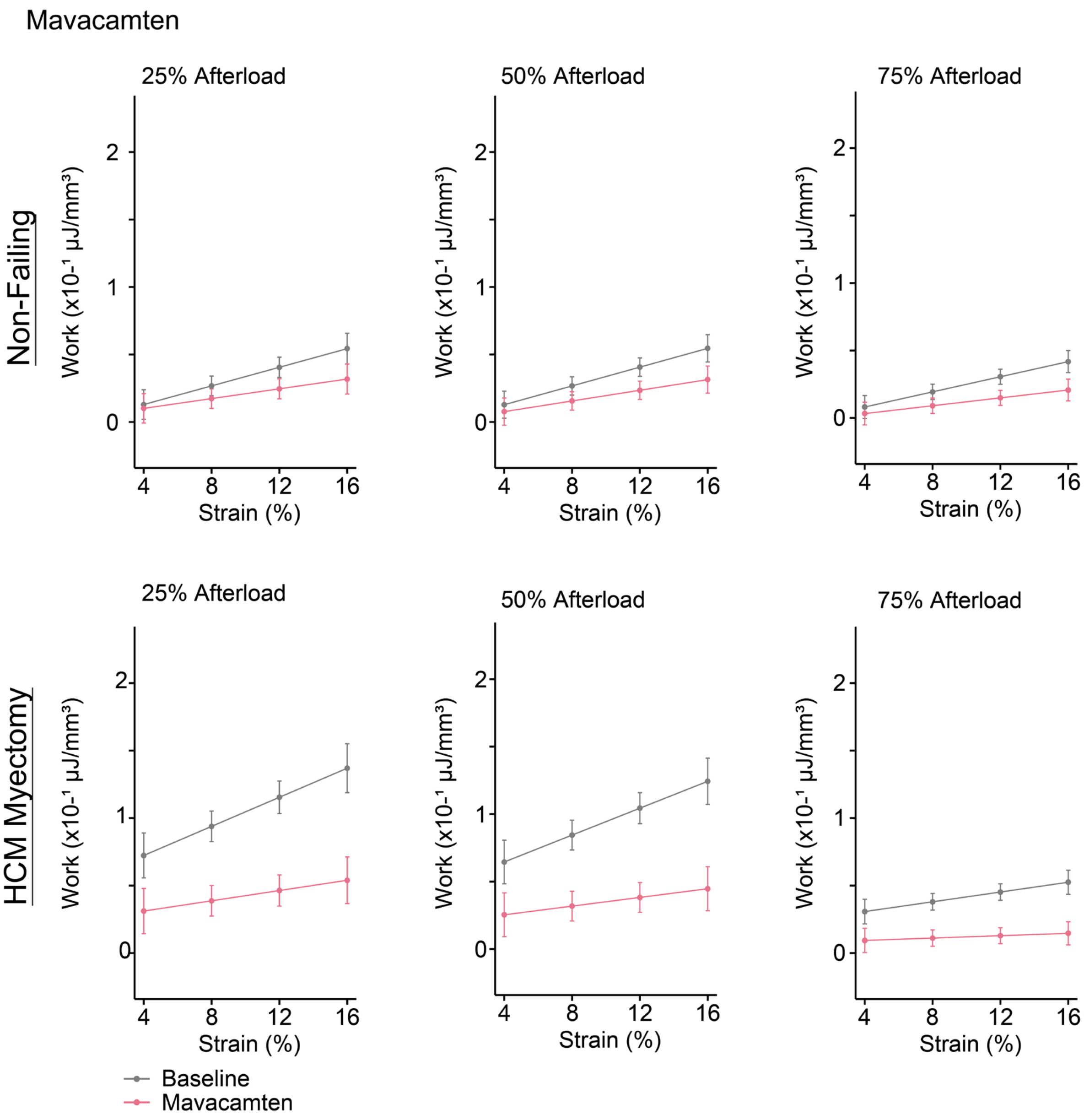

**Figure S6. Mixed Effect Linear Model Output in Response to Mavacamten.** Mean ± standard error at baseline (gray) and following mavacamten administration (pink). Replicates: N=10 hearts, n=10 (mavacamten). Statistics: Mixed-effects linear regression models with random intercepts for heart and slice

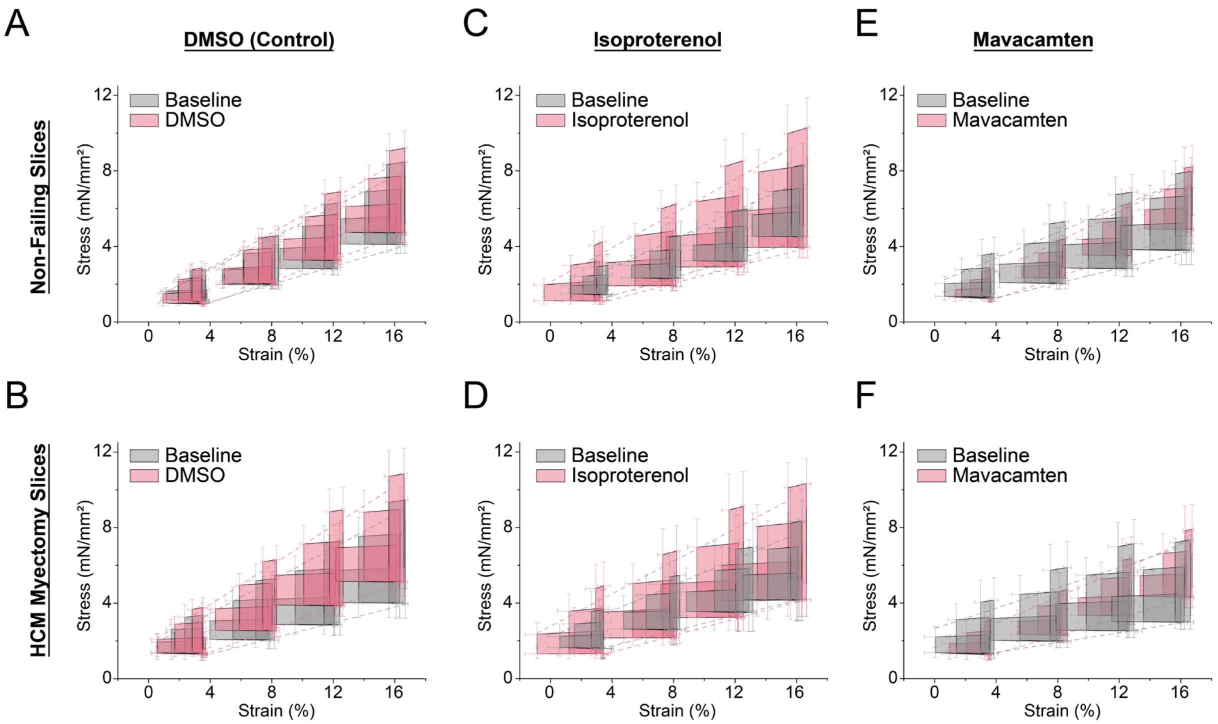

**Figure S7. Average Work Loops at Baseline and Following Drug Administration.** Average work loops ± standard error from the mean (SEM) at all preloads and afterloads. Gray work loops denote baseline measurements and pink work loops were measured following drug administration. Dotted lines refer to the end systolic and end diastolic stress strain relationships. Replicates: N=31 hearts; n=15 (DMSO), 6 (isoproterenol), 10 (mavacamten) slices.

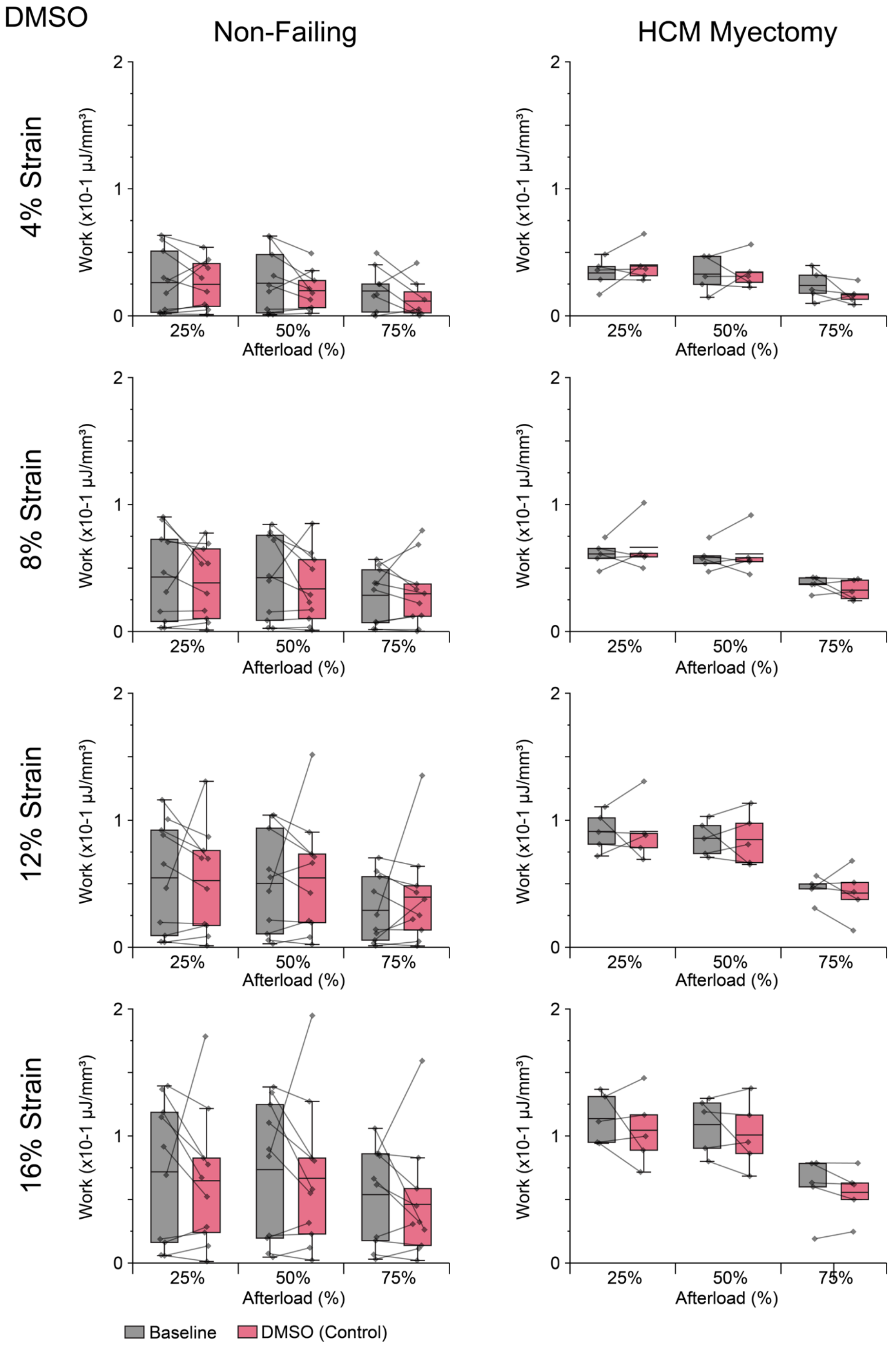

**Figure S8. Paired Measurement of Work at Baseline and Following DMSO Administration**. Each dot refers to a single measurement of work. Baseline and drug runs are paired and connected by a solid line per slice. Boxes denote median, 25$^{th}$ and 75$^{th}$ percentile and whiskers represent 1.5 interquartile range (IQR). Replicates: N=31 hearts; n=15 (DMSO), 6 (isoproterenol), 10 (mavacamten) slices.

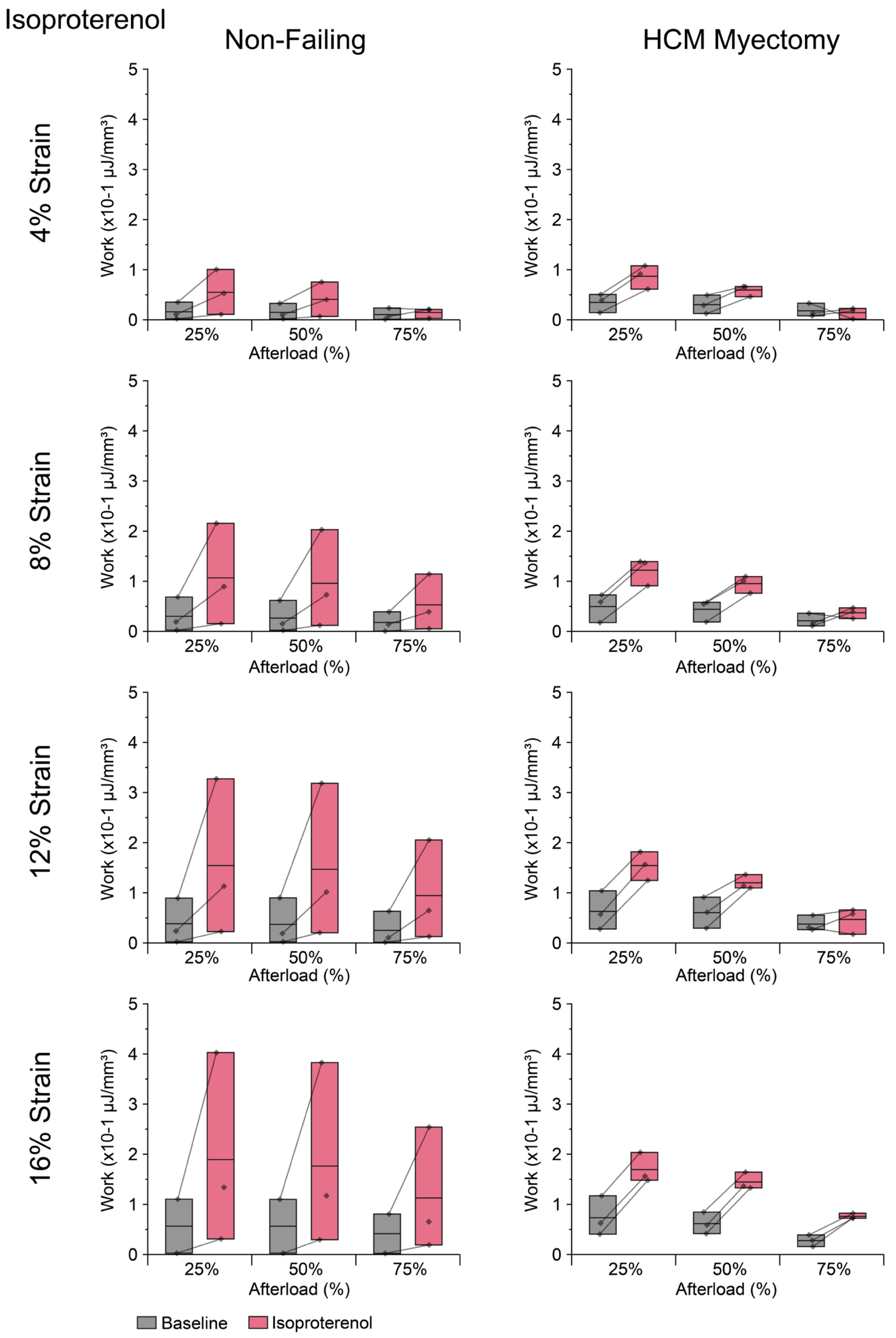

**Figure S9. Paired Measurement of Work at Baseline and Following Isoproterenol Administration**. Each dot refers to a single measurement of work. Baseline and drug runs are paired and connected by a solid line per slice. Boxes denote median, 25th and 75th percentile and whiskers represent 1.5 interquartile range (IQR). Replicates: N=6 hearts; 6 (isoproterenol), slices.

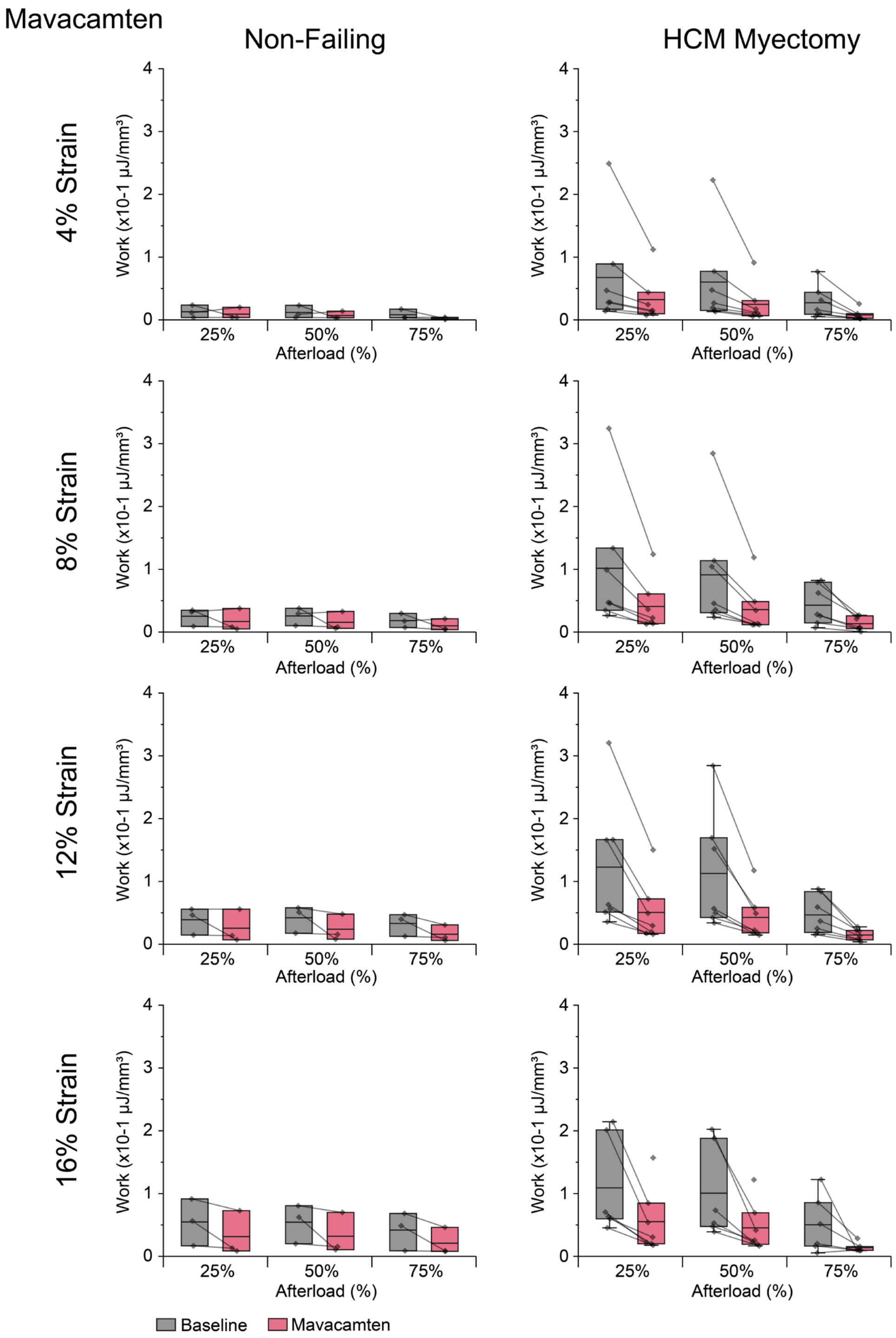

**Figure S10. Paired Measurement of Work at Baseline and Following Mavacamten Administration**. Each dot refers to a single measurement of work. Baseline and drug runs are paired and connected by a solid line per slice. Boxes denote median, 25$^{th}$ and 75$^{th}$ percentile and whiskers represent 1.5 interquartile range (IQR). Replicates: N=10 hearts; n=10 (mavacamten) slices.

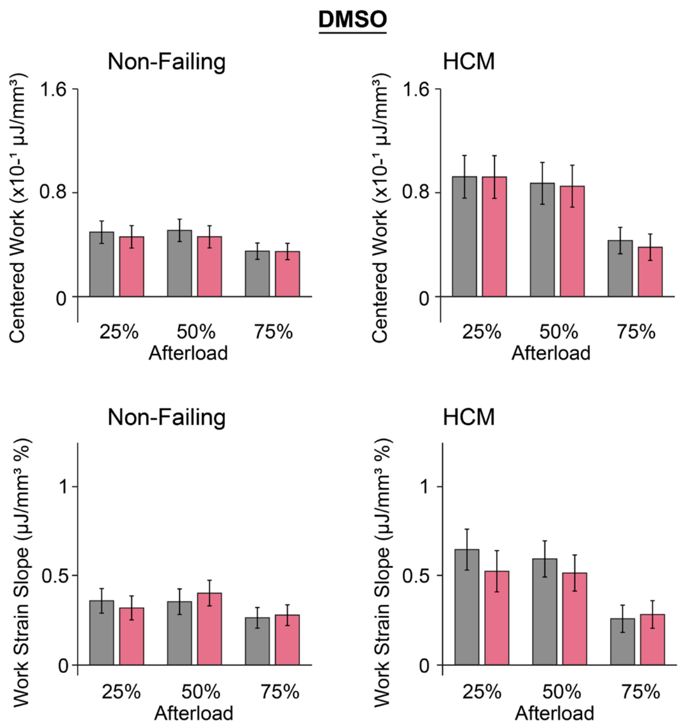

**Figure S11 Centered Work and Work-Strain Slope at Baseline and in Response to DMSO.** (Mean ± standard error). Replicates: N=15; n=15 (DMSO) slices.

| | 25% Afterload | | 50% Afterload | | 75% Afterload | |
|---|---|---|---|---|---|---|
| Condition | Dev. Stress (SEM) mN/mm$^2$ | p-value vs BL | Dev. Stress (SEM) mN/mm$^2$ | p-value vs BL | Dev. Stress (SEM) mN/mm$^2$ | p-value vs BL |
| NF | | | | | | |
| BL | 1.58 (0.15) | | 2.35 (0.24) | | 3.05 (0.33) | |
| DMSO | 1.58 (0.15) | 0.95 | 2.34 (0.24) | 0.93 | 2.93 (0.33) | 0.59 |
| BL | 1.08 (0.31) | | 1.51 (0.48) | | 2.04 (0.66) | |
| Iso | 2.93 (0.31) | **<0.0001** | 4.03 (0.48) | **<0.0001** | 5.18 (0.65) | **<0.0001** |
| BL | 0.84 (0.31) | | 1.28 (0.47) | | 1.77 (0.58) | |
| Mava | 0.59 (0.31) | 0.072 | 0.77 (0.47) | **0.016** | 1.11 (0.58) | **0.025** |
| HCM | | | | | | |
| BL | 2.23 (0.18) | | 3.31 (0.29) | | 4.35 (0.39) | |
| DMSO | 2.40 (0.18) | **0.012** | 3.57 (0.29) | **0.005** | 4.74 (0.39) | **0.002** |
| BL | 1.64 (0.25) | | 2.51 (0.40) | | 3.16 (0.54) | |
| Iso | 3.09 (0.25) | **<0.0001** | 4.19 (0.40) | **<0.0001** | 5.28 (0.54) | **<0.0001** |
| BL | 2.40 (0.26) | | 3.46 (0.41) | | 4.47 (0.56) | |
| Mava | 1.57 (0.26) | **<0.0001** | 2.25 (0.41) | **<0.0001** | 2.96 (0.56) | **<0.0001** |

**Table S3. Mean Centered Developed Active Stress (at 10% Strain) at baseline and following drug treatment**. P-values show comparison between drug treatment and baseline. P values less than 0.05 are bolded. Abbreviations: BL: baseline, Iso: isoproterenol, Mava: mavacamten. Replicates: N=31 hearts; n=15 (DMSO), 6 (isoproterenol), 10 (mavacamten) slices. Statistics: Mixed-effects linear regression models with random intercepts for heart and slice.

| | 25% Afterload | 50% Afterload | 75% Afterload |
|---|---|---|---|
| Condition | p-value vs DMSO | p-value vs DMSO | p-value vs DMSO |
| Isoproterenol | | | |
| NF, work | **<0.0001** | **<0.0001** | **0.0001** |
| HCM, work | **<0.0001** | **<0.0001** | **0.0001** |
| NF, slope | **0.016** | **0.041** | **0.044** |
| HCM, slope | **0.0001** | **<0.0001** | **0.014** |
| NF, dev st | **<0.0001** | **<0.0001** | **0.0001** |
| HCM, dev st | **<0.0001** | **<0.0001** | **0.0001** |
| Mavacamten | | | |
| NF, work | 0.32 | 0.36 | 0.13 |
| HCM, work | **<0.0001** | **<0.0001** | **<0.0001** |
| NF, slope | 0.53 | 0.34 | 0.40 |
| HCM, slope | 0.30 | 0.21 | 0.19 |
| NF, dev st | 0.12 | **0.038** | 0.13 |
| HCM, dev st | **<0.0001** | **<0.0001** | **<0.0001** |

**Table S4. Statistical Comparison Between Drug and DMSO.** For centered work at 10% strain, work-strain slope, and active developed stress at 10% strain (dev st) across all 3 afterloads. P-values show comparison between isoproterenol or mavacamten slices versus DMSO slices. P values less than 0.05 are bolded. Replicates: N=31 hearts; n=15 (DMSO), 6 (isoproterenol), 10 (mavacamten) slices. Statistics: Mixed-effects linear regression models with random intercepts for heart and slice.

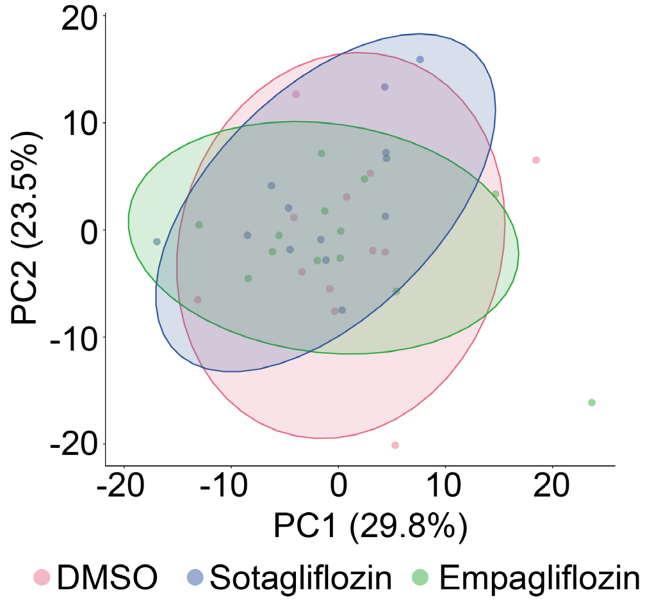

**Figure S12. PCA of the Fold Change of Work Loop Parameters in Response to SGLTi.** Each dot represents one slice and shaded circles represent 95% confidence ellipses. Replicates: N=32 hearts; n=15 (DMSO), 22 (sotagliflozin), 17 (empagliflozin) slices.

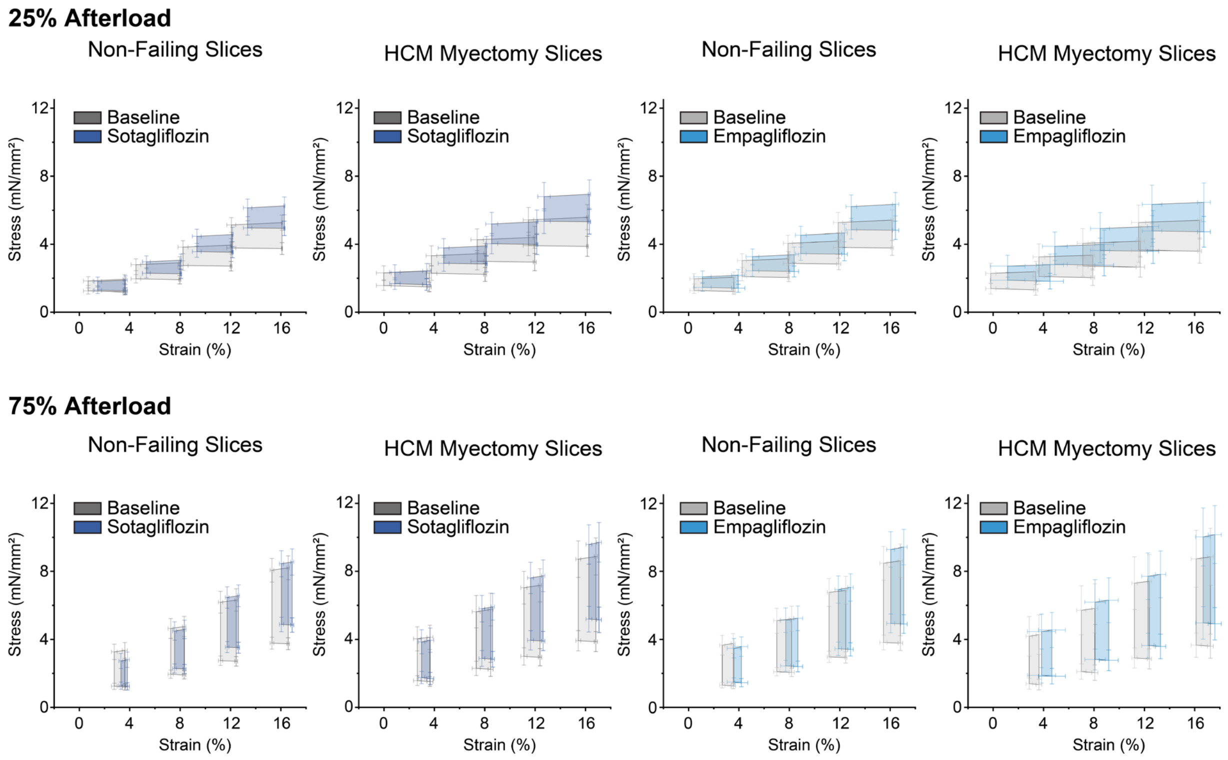

**Figure S13. Average Work Loops in Response to SGLTi**. Average work loops ± standard error from the mean (SEM) at all preloads and 25 and 75% afterloads. Replicates: N=32 hearts; n=15 (DMSO), 22 (sotagliflozin), 17 (empagliflozin) slices.

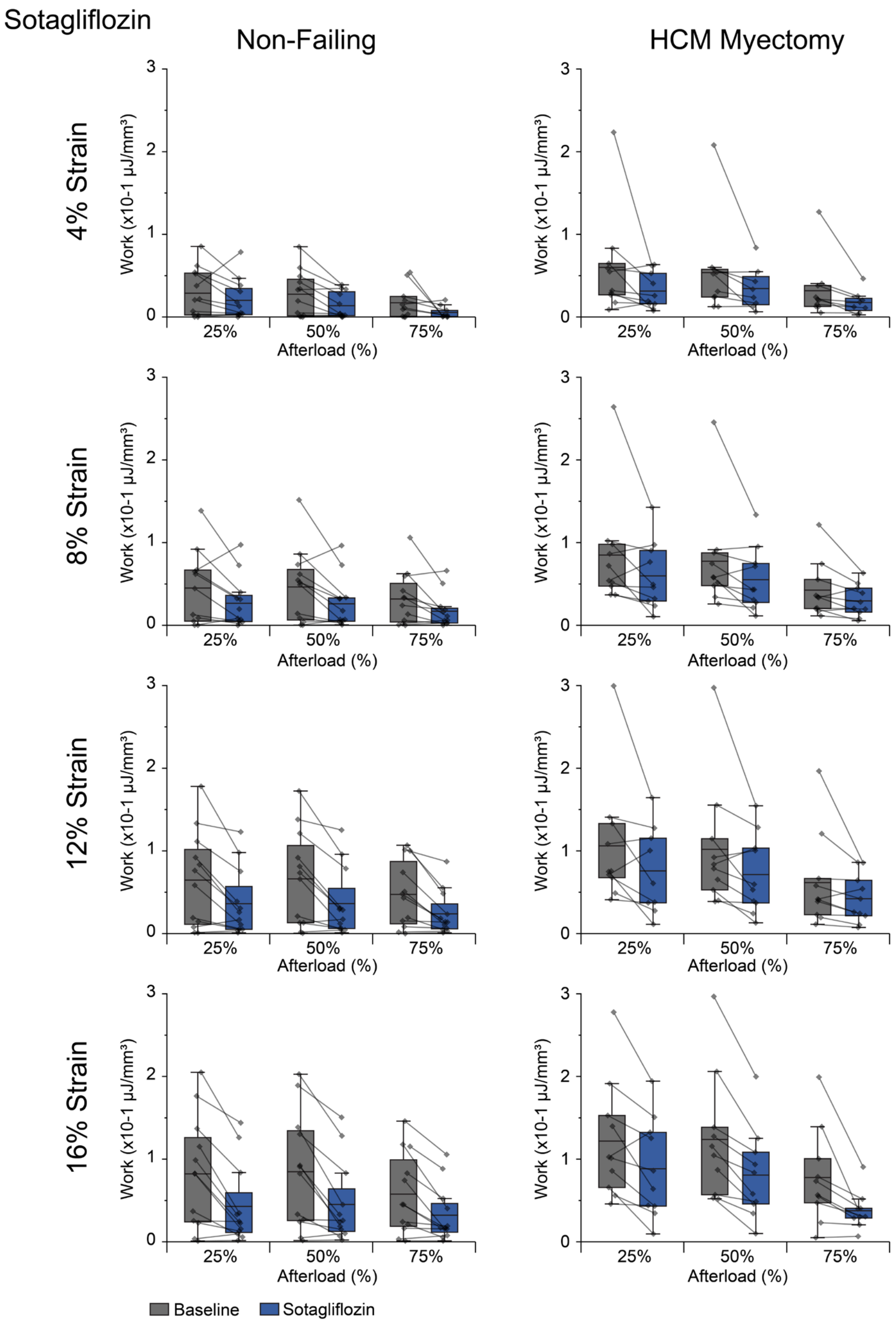

**Figure S14. Paired Measurement of Work at Baseline and Following Sotagliflozin Administration**. Each dot refers to a single measurement of work. Baseline and drug runs are paired and connected by a solid line per slice. Boxes denote median, 25th and 75th percentile and whiskers represent 1.5 interquartile range (IQR). Replicates: N=32 hearts; n=15 (DMSO), 22 (sotagliflozin) slices.

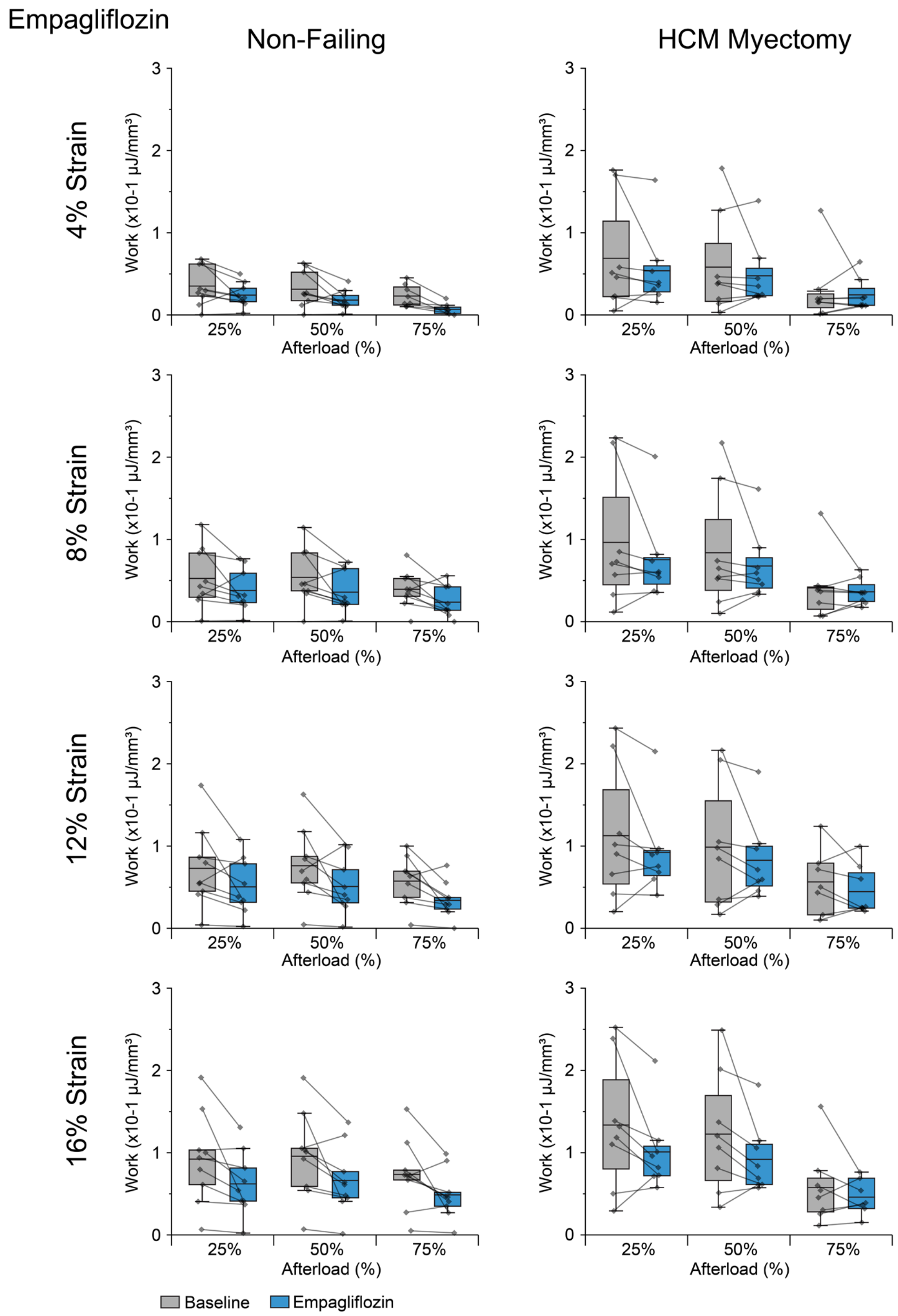

**Figure S15. Paired Measurement of Work at Baseline and Following Empagliflozin Administration**. Each dot refers to a single measurement of work. Baseline and drug runs are paired and connected by a solid line per slice. Boxes denote median, 25$^{th}$ and 75$^{th}$ percentile and whiskers represent 1.5 interquartile range (IQR). Replicates: N=32 hearts; n=15 (DMSO), 17 (empagliflozin) slices.

| | 25% Afterload | | 50% Afterload | | 75% Afterload | |
|---|---|---|---|---|---|---|
| Condition | Dev. Stress (SEM) mN/mm$^2$ | p-value vs BL | Dev. Stress (SEM) mN/mm$^2$ | p-value vs BL | Dev. Stress (SEM) mN/mm$^2$ | p-value vs BL |
| NF | | | | | | |
| BL | 1.60 (0.19) | | 2.32 (0.29) | | 3.09 (0.37) | |
| Sota | 1.21 (0.19) | **<0.0001** | 1.75 (0.29) | **<0.0001** | 2.34 (0.37) | **<0.0001** |
| BL | 1.82 (0.22) | | 2.65 (0.33) | | 3.45 (0.43) | |
| Empa | 1.53 (0.22) | **0.0006** | 2.23 (0.33) | **0.002** | 2.94 (0.43) | **0.006** |
| HCM | | | | | | |
| BL | 2.39 (0.24) | | 3.48 (0.37) | | 4.50 (0.49) | |
| Sota | 2.09 (0.24) | **<0.0001** | 3.13 (0.37) | **0.0007** | 4.00 (0.49) | **<0.0001** |
| BL | 2.40 (0.25) | | 3.38 (0.38) | | 4.30 (0.51) | |
| Empa | 2.18 (0.24) | **0.0021** | 3.08 (0.38) | **0.0085** | 3.95 (0.51) | **0.010** |

**Table S5. Mean Centered Developed Active Stress (at 10% Strain) at baseline and following SGLTi treatment**. P-values show comparison between SGLTi treatment and baseline. P values less than 0.05 are bolded. Replicates: N=32 hearts; n=15 (DMSO), 22 (sotagliflozin), 17 (empagliflozin) slices. Statistics: Mixed-effects linear regression models with random intercepts for heart and slice, with post hoc pairwise comparisons and linear combinations of model parameters. Abbreviations: BL: baseline, Sota: sotagliflozin, Empa: empagliflozin

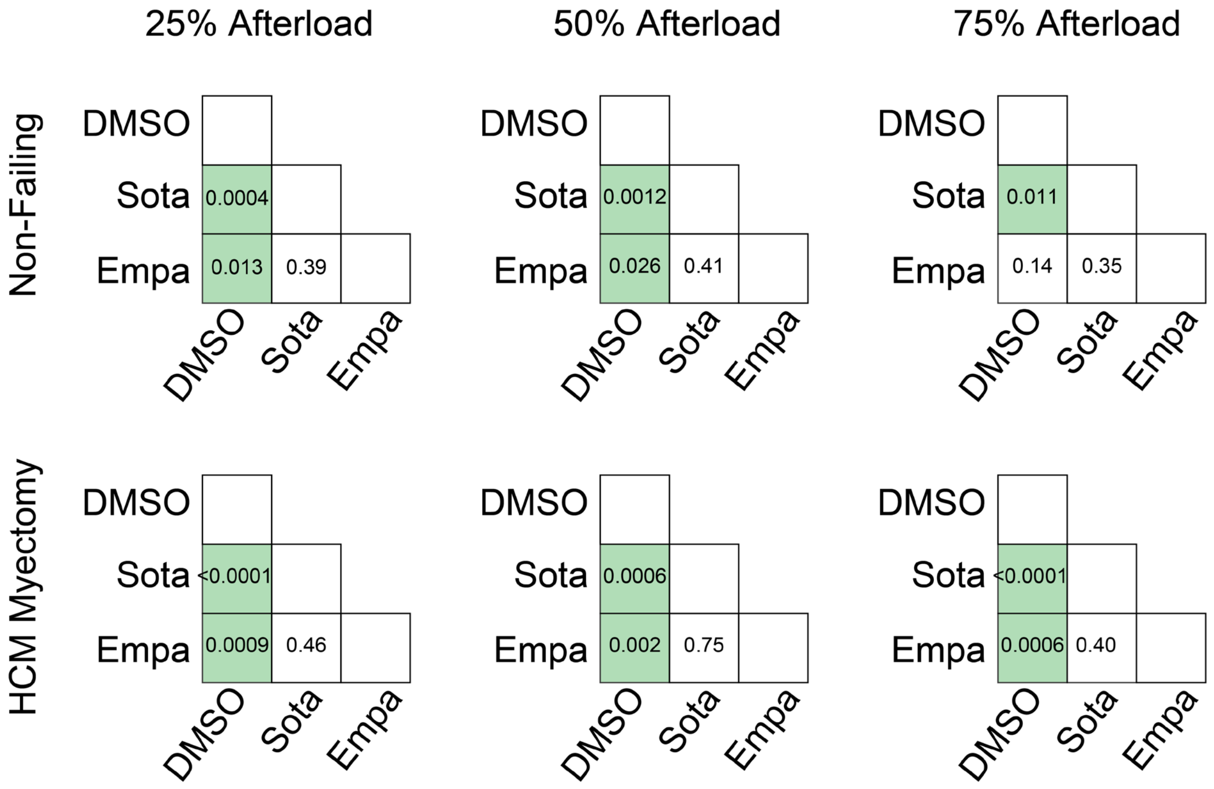

**Figure S16. Cross Drug Comparisons of Mean Developed Stress Between SGLTi and DMSO treated Slices.** Each value represents a p-value comparing SGLTis and/or DMSO. Replicates: N=32 hearts; n=15 (DMSO), 22 (sotagliflozin), 17 (empagliflozin) slices. Statistics: Mixed-effects linear regression models with random intercepts for heart and slice, with post hoc pairwise comparisons and linear combinations of model parameters.

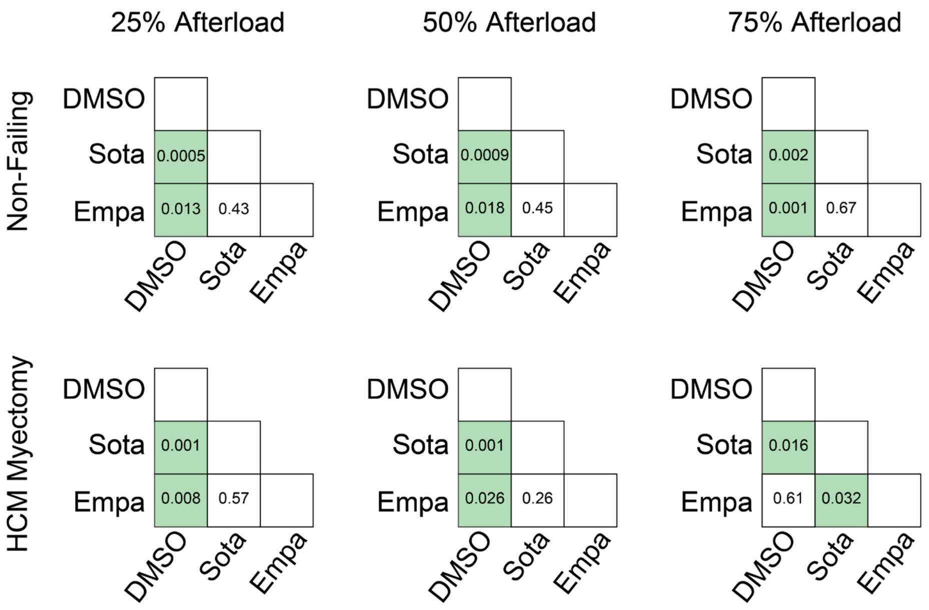

**Figure S17. Cross Drug Comparisons of Work Between SGLTi and DMSO treated Slices.** Each value represents a p-value comparing SGLTis and/or DMSO. P values less than 0.05 are shaded green. Replicates: N=32 hearts; n=15 (DMSO), 22 (sotagliflozin), 17 (empagliflozin) slices. Statistics: Mixed-effects linear regression models with random intercepts for heart and slice, with post hoc pairwise comparisons and linear combinations of model parameters.

| | 25% Afterload | | 50% Afterload | | 75% Afterload | |
|---|---|---|---|---|---|---|
| Condition | EDSSR (SEM) mN/mm²% | p-value vs BL | EDSSR (SEM) mN/mm²% | p-value vs BL | EDSSR (SEM) mN/mm²% | p-value vs BL |
| Non-Failing | | | | | | |
| BL (D) | 25.3 (2.3) | | 24.8 (2.3) | | 23.8 (2.2) | |
| DMSO | 29.6 (2.3) | 0.18 | 28.8 (2.2) | 0.21 | 30.0 (2.2) | 0.052 |
| BL (S) | 21.5 (2.1) | | 21.2 (2.0) | | 20.4 (2.0) | |
| Sota | 30.1 (2.1) | **0.0037** | 29.2 (2.0) | **0.0054** | 29.5 (2.1) | **0.0018** |
| BL (E) | 23.3 (2.5) | | 22.3 (2.3) | | 22.2 (2.3) | |
| Empa | 29.7 (2.4) | 0.062 | 28.6 (2.3) | 0.054 | 29.9 (2.3) | **0.021** |
| HCM | | | | | | |
| BL (D) | 21.3 (2.9) | | 20.6 (2.8) | | 19.8 (2.8) | |
| DMSO | 30.7 (3.0) | **0.025** | 29.6 (2.8) | **0.025** | 28.8 (2.8) | **0.024** |
| BL (S) | 18.7 (2.1) | | 17.8 (2.0) | | 17.4 (2.0) | |
| Sota | 28.7 (2.0) | **0.0006** | 27.2 (2.1) | **0.0011** | 26.6 (2.1) | **0.0016** |
| BL (E) | 17.5 (2.3) | | 16.6 (2.1) | | 16.3 (2.2) | |
| Empa | 22.1 (2.3) | 0.14 | 21.6 (2.2) | 0.106 | 21.4 (2.2) | 0.10 |

**Table S6. Mixed Linear Regression Modeling of End Diastolic Stress Strain Relationship Slope**. Values are denoted as mean (standard error). P-values show comparison between DMSO or SGLTi treated run with baseline. P values less than 0.05 are bolded. Replicates: N=32 hearts; n=15 (DMSO), 22 (sotagliflozin), 17 (empagliflozin) slices. Statistics: Mixed-effects linear regression models with random intercepts for heart and slice, with post hoc pairwise comparisons and linear combinations of model parameters.

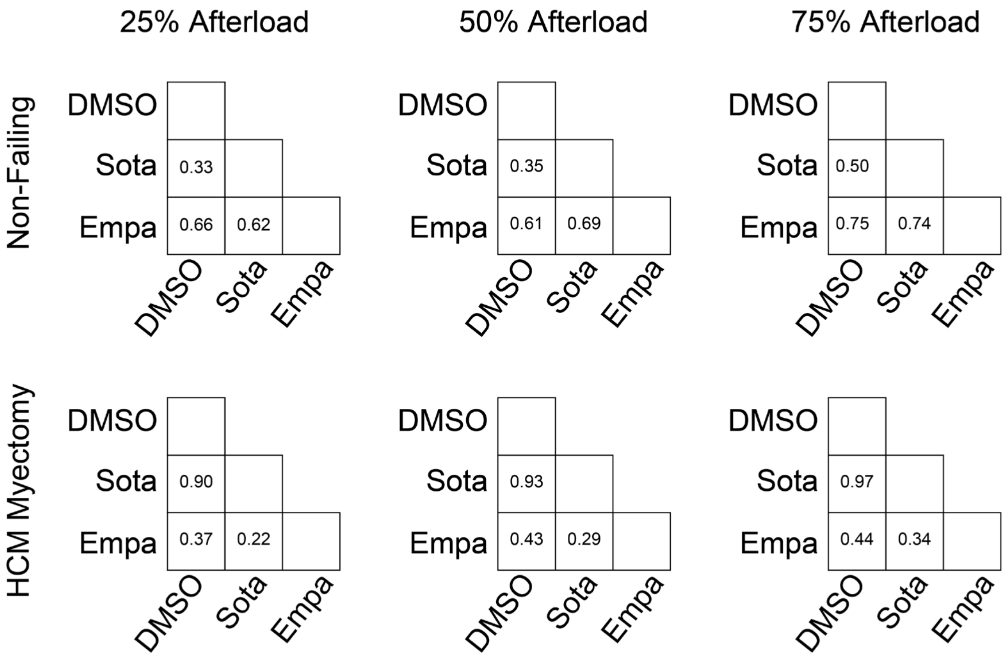

**Figure S18. Cross Drug Comparisons of End Diastolic Stress Strain Slope Between SGLTi and DMSO treated Slices.** Replicates: N=32 hearts; n=15 (DMSO), 22 (sotagliflozin), 17 (empagliflozin) slices. Statistics: Mixed-effects linear regression models with random intercepts for heart and slice, with post hoc pairwise comparisons and linear combinations of model parameters.

| | Non-Failing | | HCM | |
|---|---|---|---|---|
| Metabolite | DMSO pmol/mg (CI) | SGLTi pmol/mg (CI) | DMSO pmol/mg (CI) | SGLTi pmol/mg (CI) |
| 3-HBA | 273.8 (178, 369.59) | 488.54 (-270.86, 1247.95) | 250.88 (121.3, 380.46) | 162.71 (98.81, 226.6) |
| alpha-KG Acid | 54.07 (35.26, 72.88) | 48.71 (31.77, 65.64) | 42.45 (32.82, 52.08) | 55.72 (39.88, 71.56) |
| Citrate | 668.66 (457.27, 880.05) | 703.89 (498.91, 908.87) | 1094.68 (448.46, 1740.9) | 942.6 (681.8, 1203.4) |
| Fumarate | 23.58 (7.42, 39.73) | 31.03 (21.34, 40.73) | 15.3 (1.63, 28.96) | 33.23 (21.32, 45.15) |
| Lactate | 5537.35 (2267.94, 8806.76) | 5184.15 (4134.3, 6233.99) | 4726.49 (-550.76, 10003.74) | 4417.29 (2967.32, 5867.25) |
| Malate | 222.49 (157.4, 287.59) | 217.33 (183.24, 251.42) | 196.89 (111.6, 282.18) | 205.7 (158.91, 252.49) |
| Pyruvate | 163.34 (91.19, 235.48) | 150.14 (124.02, 176.27) | 107.45 (63.28, 151.61) | 115.97 (84.24, 147.69) |
| Succinate | 71.45 (10.42, 132.48) | 68.11 (42.89, 93.33) | 75.62 (-52.23, 203.46) | 37.85 (21.74, 53.95) |
| C02 | 1669.52 (1289.52, 2049.51) | 1728.91 (1505.56, 1952.26) | 1238.37 (502.59, 1974.15) | 1441.39 (1068.34, 1814.45) |
| C03 | 9.64 (4.75, 14.52) | 9.96 (7.06, 12.87) | 4.32 (1.64, 6.99) | 6.07 (3.69, 8.44) |
| C03-DC | 0.67 (0.18, 1.17) | 0.86 (0.57, 1.15) | 0.41 (0.29, 0.53) | 0.96 (0.58, 1.33) |
| C04 Butyryl | 7.68 (3.44, 11.92) | 7.61 (4.77, 10.45) | 8.12 (-2.81, 19.06) | 5.78 (1.09, 10.48) |
| C04 Isobutyryl | 2.14 (0.67, 3.61) | 3.46 (1.42, 5.51) | 9.79 (-5.39, 24.97) | 2.52 (0.31, 4.73) |
| C04-DC MeMal | 0.46 (0.33, 0.59) | 1.25 (-0.21, 2.71) | 0.4 (0.28, 0.52) | 0.43 (0.32, 0.53) |
| C04-DC Succinyl | 1.52 (0.37, 2.67) | 2.97 (0.62, 5.33) | 0.62 (0.24, 1.01) | 0.64 (0.41, 0.87) |
| C04-OH Butyryl | 11.57 (6.28, 16.86) | 15.91 (10.69, 21.14) | 7.49 (2.32, 12.67) | 7.74 (4.35, 11.13) |
| C04-OH Isobutyryl | 56.73 (17.85, 95.6) | 50.69 (31.01, 70.38) | 24.18 (-2.4, 50.75) | 26.07 (8.03, 44.12) |
| C05 2-Methylbutyryl | 2.74 (0.67, 4.8) | 3.58 (2.31, 4.84) | 1.26 (0.64, 1.89) | 1.08 (0.89, 1.28) |
| C05 Isovaleryl | 1.94 (1.28, 2.6) | 2.38 (1.65, 3.11) | 2.55 (-0.48, 5.58) | 1.31 (0.87, 1.75) |
| C05:1 | 3.87 (1.21, 6.53) | 4.32 (2.53, 6.1) | 1.92 (1.31, 2.53) | 1.4 (0.91, 1.89) |
| C05-DC | 0.62 (0.31, 0.93) | 0.69 (0.43, 0.95) | 0.43 (0.2, 0.67) | 0.54 (0.36, 0.71) |
| C05-OH | 15.56 (3.87, 27.25) | 14.23 (6.11, 22.34) | 8.5 (4.93, 12.07) | 5.76 (4.17, 7.34) |
| C06 | 1.17 (0.49, 1.85) | 1.04 (0.61, 1.48) | 0.61 (0.1, 1.12) | 0.61 (0.39, 0.84) |
| C06-OH | 1.24 (0.94, 1.55) | 1.59 (1.14, 2.05) | 1.66 (0.88, 2.44) | 1.33 (0.83, 1.84) |
| C08 | 0.42 (0.07, 0.77) | 0.43 (0.23, 0.64) | 0.25 (0.09, 0.41) | 0.28 (0.14, 0.42) |
| C08:1-OH | 0.18 (0.03, 0.33) | 0.27 (0.13, 0.41) | 0.57 (0.36, 0.78) | 0.43 (0.26, 0.6) |
| C08-OH | 0.71 (0.26, 1.16) | 0.69 (0.38, 0.99) | 0.23 (-0.02, 0.48) | 0.33 (0.18, 0.49) |
| C10 | 1.86 (-0.25, 3.97) | 2.71 (1.69, 3.72) | 0.8 (-0.62, 2.22) | 2.89 (1.66, 4.12) |
| C10_div10run2 | 0.27 (0.1, 0.43) | 0.38 (0.31, 0.44) | 0.26 (0.14, 0.39) | 0.41 (0.33, 0.48) |
| C10-OH | 0.47 (-0.09, 1.02) | 0.43 (0.22, 0.64) | 0.07 (0, 0.14) | 0.23 (0.11, 0.34) |
| C12 | 2.81 (0.76, 4.86) | 2.45 (1.43, 3.47) | 2.69 (0.84, 4.54) | 2.65 (1.44, 3.86) |
| C12:1 | 0.86 (0.2, 1.53) | 0.78 (0.48, 1.08) | 0.5 (0.27, 0.73) | 0.6 (0.4, 0.8) |
| C12-OH | 1.11 (0.29, 1.93) | 0.91 (0.63, 1.19) | 0.51 (0.22, 0.79) | 0.67 (0.44, 0.91) |

| | | | | |
|---|---|---|---|---|
| C14 | 4.49 (-0.16, 9.14) | 3.89 (2.33, 5.44) | 6.46 (1.42, 11.49) | 6.15 (2.23, 10.07) |
| C14:1 | 3.39 (-0.23, 7.01) | 2.7 (1.39, 4) | 2.23 (0.5, 3.95) | 2.39 (1.47, 3.3) |
| C14:1-OH | 0.4 (0.09, 0.72) | 0.41 (0.19, 0.63) | 0.17 (0.06, 0.29) | 0.23 (0.15, 0.31) |
| C14:2 | 1.1 (0.43, 1.76) | 0.76 (0.49, 1.03) | 0.83 (0.3, 1.36) | 0.91 (0.51, 1.32) |
| C14:2-OH | 0.12 (0.08, 0.15) | 0.1 (0.06, 0.15) | 0.13 (0.07, 0.2) | 0.09 (0.05, 0.13) |
| C14-OH | 1.03 (0.26, 1.8) | 1.06 (0.69, 1.42) | 0.8 (0.32, 1.29) | 0.84 (0.47, 1.21) |
| C16 | 19.26 (0.99, 37.53) | 15.61 (7.56, 23.66) | 32.23 (7.56, 56.9) | 29.92 (11.8, 48.04) |
| C16:1 | 7.03 (-1.91, 15.97) | 5.92 (2.48, 9.36) | 10.61 (0.83, 20.4) | 8.3 (3.11, 13.49) |
| C16:1-OH | 1.33 (0.01, 2.65) | 1.25 (0.82, 1.68) | 1.28 (0.38, 2.19) | 1.19 (0.6, 1.77) |
| C16:2 | 0.91 (0.09, 1.73) | 0.97 (0.46, 1.48) | 3.2 (0.35, 6.05) | 2.27 (0.7, 3.85) |
| C16:2-OH | 0.37 (0.16, 0.58) | 0.37 (0.27, 0.48) | 0.33 (0.13, 0.53) | 0.39 (0.24, 0.54) |
| C16-OH | 2.01 (0.2, 3.81) | 2.06 (1.24, 2.89) | 3.64 (0.52, 6.77) | 2.96 (1.18, 4.73) |
| C18 | 14.64 (-0.86, 30.14) | 13.97 (7.54, 20.39) | 18.12 (3.92, 32.33) | 23.7 (9.99, 37.41) |
| C18:1 | 83.64 (-51.13, 218.41) | 54.68 (28.38, 80.98) | 73.65 (6.19, 141.12) | 102.66 (30.15, 175.16) |
| C18:1-OH | 7.01 (-2.41, 16.42) | 5.9 (3.58, 8.21) | 6.86 (0.9, 12.81) | 8.11 (2.5, 13.73) |
| C18:2 | 25.38 (-9.52, 60.28) | 18.58 (11.26, 25.91) | 34.47 (5.21, 63.73) | 53.66 (13.08, 94.24) |
| C18:2-OH | 1.5 (-0.3, 3.3) | 1.73 (1.09, 2.36) | 3.01 (0.59, 5.43) | 3.53 (0.49, 6.56) |
| C18-OH | 1.88 (-0.6, 4.36) | 1.65 (0.9, 2.41) | 2.35 (0.3, 4.4) | 2.84 (0.56, 5.13) |
| C20 | 0.78 (-0.27, 1.83) | 1.06 (0.17, 1.95) | 1.99 (-0.21, 4.19) | 1.8 (0.53, 3.08) |

**Table S7. Targeted Metabolomics of LMS following DMSO or SGLTi.** Mean amount of select metabolites (pmol) relative to tissue dry mass with 95% Confidence Intervals (CI). In the SGLTi group, both sotagliflozin and empagliflozin slices were included. Replicates: N=32 hearts; n=8 (NF, DMSO), 15 (NF, SGLTi), 6 (HCM, DMSO), 14 (HCM, SGLTi) slices.

### REFERENCES

1. Chen CY, Caporizzo MA, Bedi K, Vite A, Bogush AI, Robison P, Heffler JG, Salomon AK, Kelly NA, Babu A, et al. Suppression of detyrosinated microtubules improves cardiomyocyte function in human heart failure. *Nat Med*. 2018;24:1225–1233. doi: 10.1038/s41591-018-0046-2
2. Lanfear DE, Gibbs JJ, Li J, She R, Petucci C, Culver JA, Tang WHW, Pinto YM, Williams LK, Sabbah HN, et al. Targeted Metabolomic Profiling of Plasma and Survival in Heart Failure Patients. *JACC Heart failure*. 2017;5:823–832. doi: 10.1016/j.jchf.2017.07.009
